# Targeting *de novo* lipogenesis improves gemcitabine efficacy in pancreatic ductal adenocarcinoma

**DOI:** 10.1101/2024.12.18.628624

**Authors:** Sarah E Hancock, Linda Garthwaite, Konstantina Harellis, Michael Susetio, Eileen Ding, Laura Choong, Osvaldo Contreras, Amy Nguyen, Josiah Lising, Felicia KM Hansen, Puttandon Wongsomboon, Jan Philipp Menzel, Berwyck LJ. Poad, Todd W Mitchell, Stephen J Blanksby, Nigel Turner

## Abstract

Pancreatic ductal adenocarcinoma (PDAC) is a highly aggressive disease with few treatment options and poor survivability. In this work we sought to characterise metabolic adaptations to gemcitabine (GEMC)-based chemotherapy exposure to discover new therapeutic targets for improving treatment efficacy. We show that GEMC resistance (GEMR) upregulates *de novo* lipogenesis in Panc1 and MiaPaCa2 cells through increased activity and expression of acetyl-CoA carboxylase (ACC), fatty acid synthase (FAS) and stearoyl-CoA desaturase 1 (SCD1). We also discovered alternate fatty acid desaturase 2 (FADS2) activity in Panc1 cells, which led to the production of sapienic acid (FA 16:1*n*-10, *cis*) from palmitic acid (FA 16:0). Knockdown of key lipid synthesis enzymes sensitised cells to GEMC treatment, with FAS (both cell lines), SCD1 (MiaPaCa2 only) and SCD1+FADS2 (Panc1) knockdown showing the greatest reduction in cell growth when combined with GEMC treatment. In Panc1 cells, both desaturases upregulated their activity when the alternate was knocked down, necessitating the need for dual desaturase knockdown in this cell line. PDAC cells attenuated to grow in combination GEMC/paclitaxel (CombAT) also displayed enhanced *de novo* lipogenesis; however, combination chemotherapy significantly downregulated FADS2 expression and activity in Panc1 CombAT cells rendering them more sensitive to SCD1 knockdown. We conclude that co-targeting lipid synthesis in PDAC could be a viable strategy for improving the efficacy of both GEMC monotherapy and combination GEMC/PTX therapy.

## Introduction

Pancreatic ductal adenocarcinoma (PDAC) is the twelfth most common cancer worldwide, with close to half a million new cases being diagnosed in 2020 ^1^. Despite its relatively low incidence rate, the mortality associated with PDAC remains high, with a global 5-year overall survival rate of just 9% ^2^. The ongoing poor survivability of PDAC is a result of many contributing factors. PDAC is a highly asymptomatic disease that is often detected at later stages, with fewer than 20% of cases being eligible for surgical resection at the time of diagnosis ^3^. For most patients only palliative chemotherapy remains as a treatment option. Either FOLFIRINOX (folinic acid, 5-fluorouracil, irinotecan and oxaliplatin) or gemcitabine (GEMC) monotherapy have long been the primary chemotherapy options for PDAC, with the former coming at a cost of higher toxicity ^4,5^. More recently, GEMC combined with albumin*-*bound (nab-)paclitaxel (also known as Abraxane) has replaced GEMC monotherapy as the standard of care for metastatic PDAC, but survivability still remains extremely poor with an increase in median overall survival of only a few months when compared with GEMC monotherapy ^6^.

Underlying the poor treatability and chemoresistance of PDAC are a myriad of genetic mutations, many of which are sporadic and heterogeneous between patients. Activating mutations in *KRAS* are common and present in around 90% of tumours, as are inactivating mutations in *TP53*, *CDKN2A*, and *SMAD4* (>20% of tumours) ^3,7^. Beyond this, common mutations seen across PDAC tumours drop to <10% ^3,8^, precluding the identification of druggable gene therapy targets that can impact a majority of patients. The genetic mutations present in PDAC have wide-ranging impacts on tumour molecular biology, activating diverse pro-survival and anti-apoptotic signalling pathways ^9–11^. Further compounding the chemoresistant nature of PDAC is the presence of a dense desmoplastic stroma that creates a hypoxic tumour microenvironment and impedes the delivery of chemotherapy to cancer cells ^12^. Together this results in extensive metabolic reprogramming in PDAC to facilitate the acquisition of biomolecules necessary to support cancer cell rapid growth and proliferation. Despite the lack of a good genetic therapeutic target in PDAC, mutational effects likely converge on relatively fewer metabolic pathways ^13,14^, making metabolism-based therapies an attractive pathway for improving options for PDAC treatment.

The role of lipid metabolism in cancer cell survival, metastasis, and chemoresistance has become a recent topic of interest across many different cancer types including PDAC ^15–21^. Key *de novo* lipogenesis enzymes are commonly upregulated in many cancers, consistent with the continual need for accrual of lipids in cancer ^22^. For example, fatty acid synthase (FAS) is the rate-limiting step in fatty acid synthesis and catalyses the elongation of malonyl-CoA produced by acetyl-CoA carboxylase (ACC) to palmitic acid (*i.e.*, FA 16:0, a fatty acid with 16 carbons and no carbon*-*carbon double bonds). FAS is commonly expressed at high levels in a variety of cancers (reviewed by ^23–25)^, where its overexpression often correlates to poorer survival ^17,26^ and chemotherapy resistance ^17,27–29^. Aberrant lipid metabolism is an emerging metabolic hallmark of PDAC, where fatty acid and sterol metabolism is upregulated ^18,30^, and distinct lipid signatures can predict patient prognosis and resistance to chemotherapy ^15^. Collectively, relatively little is known about the role of lipid metabolism in PDAC growth, metastasis, and chemoresistance, and a more complete elucidation of implicated metabolic pathways could lead to the identification of novel therapeutic targets to improve PDAC treatment and patient survival.

In this work, we describe significant increases in *de novo* lipogenesis following chronic treatment of PDAC cells with GEMC monotherapy or combination GEMC/paclitaxel (PTX) therapy. We show that targeting key lipogenic enzymes can improve the efficacy of GEMC chemotherapy, including ACC1, FAS, stearoyl-CoA desaturase 1 (SCD1) and fatty acid desaturase 2 (FADS2). Furthermore, we describe Δ6 desaturation of saturated fatty acids by FADS2 in PDAC for the first time, alongside selective downregulation of FADS2 by chronic GEMC/PTX treatment offering a novel pathway towards dual desaturase targeting to improve GEMC-based chemotherapy efficacy in PDAC.

## Results

### Development of GEMR PDAC cell lines

Panc1 and MiaPaCa2 were chosen as model cell lines to explore chemotherapy-induced metabolic reprogramming due to their divergent mutational heterogeneity (Figure S1 & Table S1) combined with discrete metabolic phenotypes ^31^. GEMR cells lines from Panc1 and MiaPaCa2 cells were developed as described in Materials and methods, with the final concentration of GEMC for Panc1 cells being 150 nM and 64 nM for MiaPaCa2 cells. At these doses, Panc1 GEMR cells had an IC_50_ that was >2.5 times that of CON cells (Table 1), and MiaPaCa2 GEMR IC_50_ was ∼3 times that of CON. Few overt morphological changes could be observed between GEMR and CON in either cell line (Figure S2). Together these data suggest that both PDAC cell lines are newly adapted to GEMC, lacking the extreme changes in cell morphology seen in more advanced GEMR PDAC models ^32–34^.

**Table 1:** half-maximal inhibitory concentration of control and gemcitabine-resistant (GEMR) PDAC cell lines.

| <u>Cell line</u> | <u>$IC_{50}$ (nM)</u> | <u>95% CI (nM)</u> | <u>p value*</u> |
| --- | --- | --- | --- |
| Panc1 CON | 273.3 | 136.0 - 575.5 | - |
| Panc1 GEMR | 720.5 | 427.2 - 1225.0 | 0.012 |
| MiaPaCa2 CON | 30.6 | 20.3 - 46.1 | - |
| MiaPaCa2 GEMR | 92.2 | 59.7 - 142.2 | <0.001 |

CON control GEMR gemcitabine resistant $IC_{50}$ half-maximal inhibitory concentration.
\* control vs GEMR, non-linear regression with extra sum-of-squares F test, n= 3.

### GEMR alters lipid profile and increases lipid droplet number in Panc1 cells

Previous work had indicated a distinct lipidomic signature associated with chemoresistant PDAC ^15^. To better understand impact of GEMR on PDAC cell lipid metabolism we performed untargeted lipidomics. Untargeted lipidomics enabled the quantification of more than 400 distinct molecular lipid species within Panc1 & MiaPaCa2 control & GEMR cells from 12 different classes of lipid, with 77 molecular species being differentially abundant between Panc1 control and GEMR cells (Figure 1A) and 79 between MiaPaCa2 cell lines (Figure 1B). Roughly equal proportions of increased and decreased lipid species were detected between Panc1 CON & GEMR cells (Figure 1A), while lipid species in MiaPaCa2 GEMR cells were mostly decreased with GEMR (Figure 1B). Interestingly, Panc1 cells appeared to have an overrepresentation of lipids containing saturated (no carbon*-*carbon double bonds; SFA) or monounsaturated fatty acids (1 carbon*-*carbon double bond; MUFA) that were increased with GEMR (Figure 1A), strongly suggesting the upregulation of *de novo* lipid synthesis within this cell line. Confocal imaging of BODIPY-stained cells revealed increased lipid droplet number per cell in Panc1 GEMR cells when compared with CON, with no change in lipid droplet size (Figure 1D). No significant change in lipid droplet number or size was observed between MiaPaCa2 CON and GEMR cells (Figure 1E); however, MiaPaCa2 CON cells contained approximately three times more neutral lipid droplets per cell compared with Panc1 CON (Figure 1F). Together these data suggest a role for GEMC-driven upregulation of *de novo* lipogenesis and lipid droplet biogenesis in Panc1 cells specifically.

**Figure 1:**
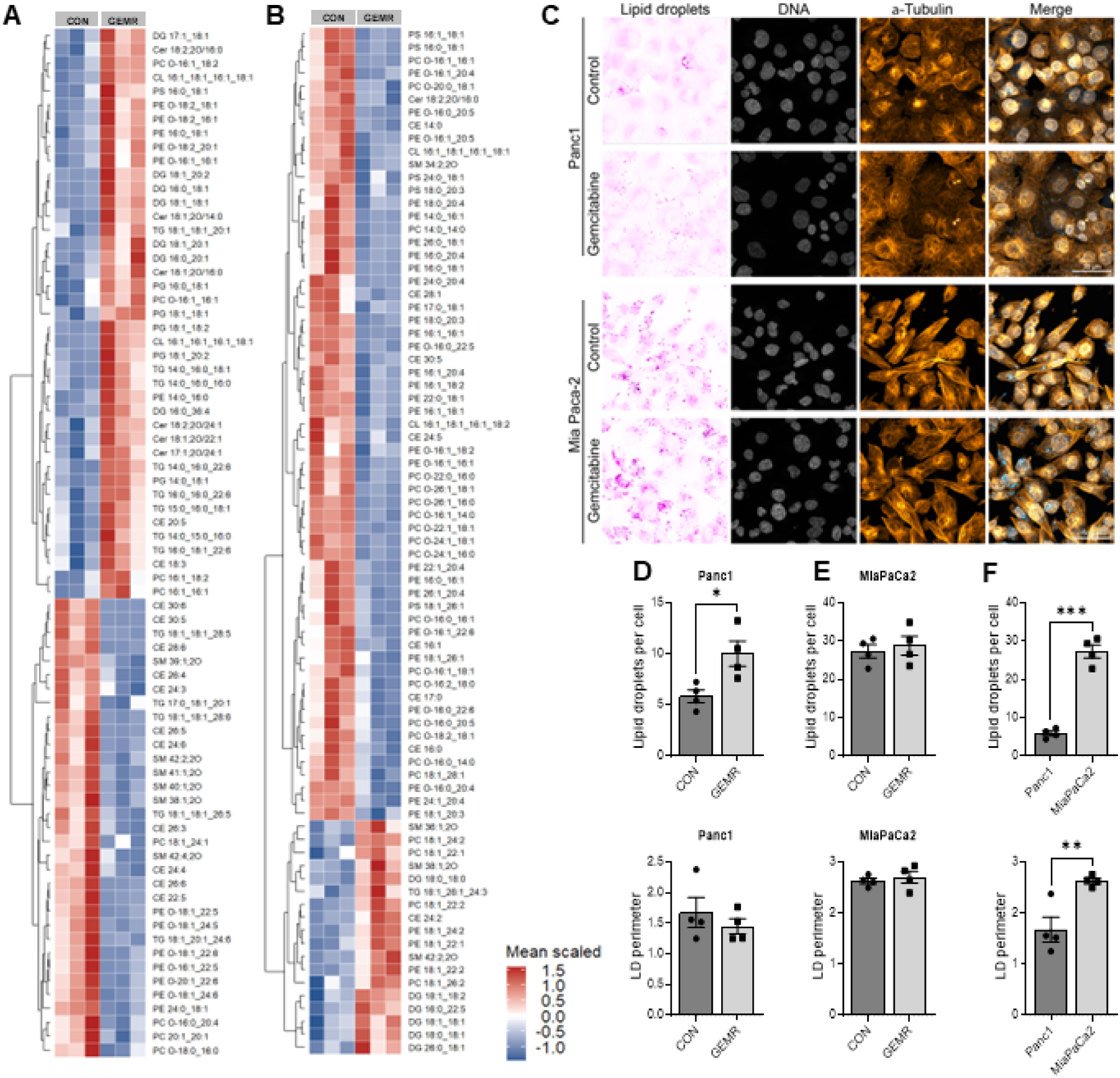
Differentially abundant lipid species from **A** Panc1 and **B** MiaPaCa2 control (CON) & gemcitabine-resitant (GEMR) cell lines detected by untargeted lipidomics (unpaired t test, *P* < 0.05). Lipids were quantified from internal standards, and values are mean*-*scaled. **C** Representative confocal images of BOPIDY-stained Panc1 & MiaPaCa2 CON & GEMR cells lines, with (**D-E**) quantification of lipid droplets per cell and lipid droplet perimeter (µm). **F** Comparison of lipid droplet number and size between Panc1 and MiaPaCa2 cells. Statistical significance was determined by unpaired t test, \**P* < 0.05 ** *P* < 0.01. *** *P* < 0.001. *CE* cholesteryl ester, *Cer* ceramide, *CL* cardiolipin, *DG* diacylglycerol, *LPC* lysophosphatidylcholine, *LPE* lysophosphatidylethanolamine, *PC* phosphatidylcholine, *PE* phosphatidylethanolamine, *PG* phosphatidylglycerol, *PS* phosphatidylserine, *SM* sphingomyelin*, TG* triacylglycerol

### GEMR increases *de novo* lipogenesis in PDAC cell lines

To better understand GEMR-driven reprogramming of PDAC lipid metabolism we undertook extensive profiling of lipid synthesis pathways in both cell lines (Figure 2A). We measured *de novo* lipogenesis in PDAC cells by tracing the incorporation of radiolabelled glucose into the lipid fraction of both Panc1 and MiaPaCa2 CON & GEMR cells (Figure 2B). Surprisingly GEMR increased *de novo* synthesis of lipid from glucose in both PDAC cell lines. To further elucidate specific changes in lipid metabolism we next performed immunoblotting to characterise changes in total and activated enzymes involved in lipid synthesis. Phosphorylation of acetyl-CoA carboxylase 1 (ACC1) at serine 79 was decreased in Panc1 GEMR (indicating enhanced activity of this enzyme) alongside an increased abundance of fatty acid synthase (FAS, Figure 2C). MiaPaCa2 GEMR cells had increased total ACC compared to CON (Figure 2D). Together, these data suggest that GEMR upregulates *de novo* lipogenesis in both PDAC cell lines.

**Figure 2:**
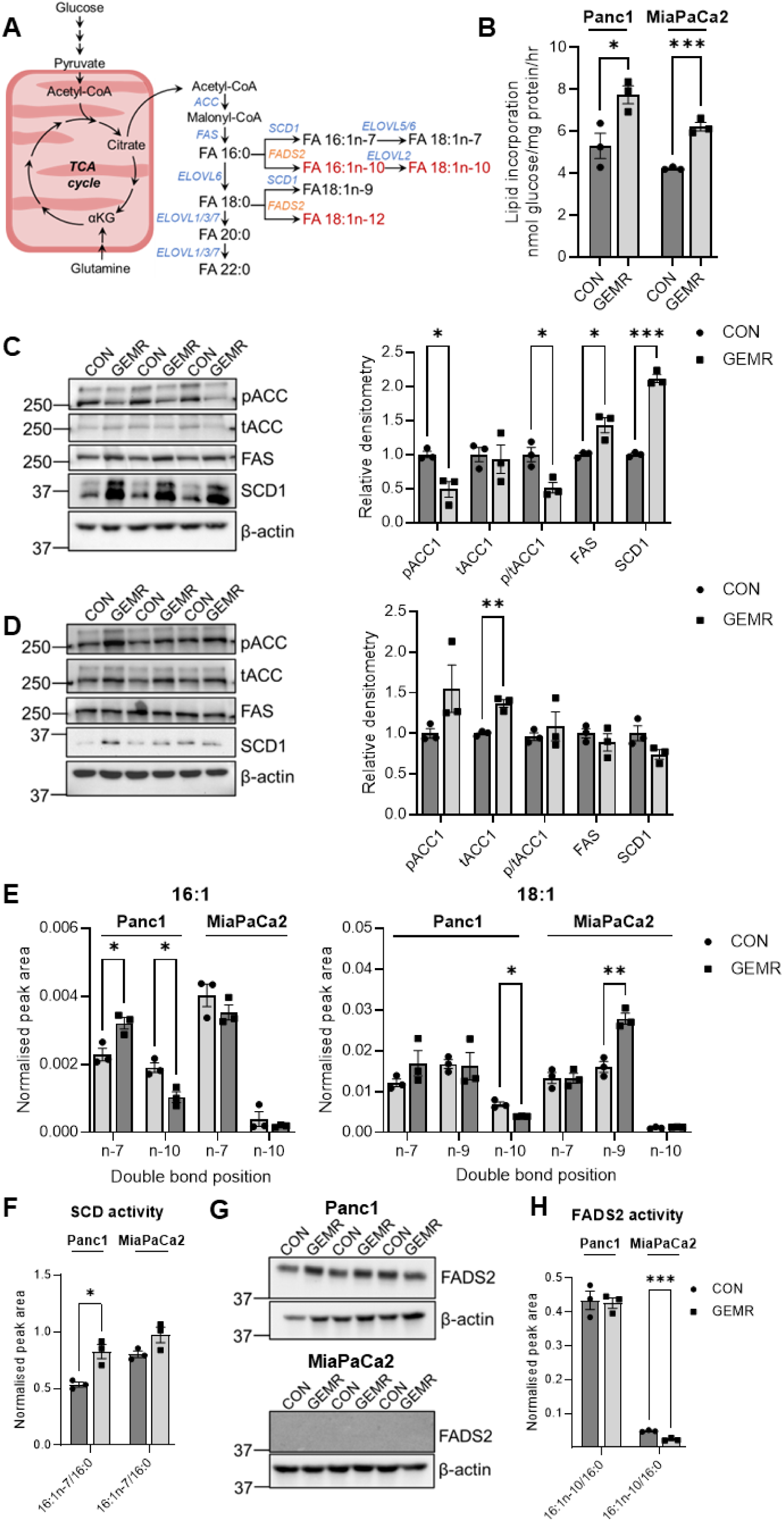
**A** map of lipid synthesis in mammalian cells, including elongation and desaturation steps. Canonical lipid synthesis pathways are shown in blue/black, while the alternative fatty acid desaturase 2 (FADS2)-mediated pathway is shown in red/orange. **B** *De novo* lipogenesis measured by tracing uniformly radiolabelled glucose into the lipid fraction of control (CON) and gemcitabine resistant (GEMR) Panc1 and MiaPaCa2 cells. Immunoblotting of key enzymes involved in *de novo* lipid synthesis in **C** Panc1 and **D** MiaPaCa2 CON and GEMR cell lines. **E** Analysis of AMP^+^-derivatised isomeric FA16:1 and FA 18:1 MUFA by ozone-enabled fatty acid discovery (OzFAD). **F** SCD1 activity measured by tracing uniformly labelled ^13^C-palmitate (FA 16:0) into palmitoleic acid (FA 16:1*n-*7) using liquid chromatography-mass spectrometry (LC-MS). **G** immunoblotting of FADS2. **H** FADS2 activity measured by tracing uniformly labelled ^13^C-palmitic acid into sapienic acid (FA 16:1*n-*10) by LC-MS. Unpaired t test, * *P* < 0.05, ** *P* < 0.01, *** *P* < 0.001. *ELOVL* elongation of very long chain fatty acids *FADS2* fatty acid desaturase *FAS* fatty acid synthase 2 *pACC* phosphorylated acetyl-CoA carboxylase *SCD1* stearoyl-CoA desaturase 1 *tACC* total ACC.

Given the upregulation of lipid synthesis with GEMR in both cell lines we also assessed the impact of GEMR on fatty acid oxidation (FAO) using radiolabelled palmitate. We found increased FAO in Panc1 GEMR cell lines and minimal amounts of FAO in both CON and GEMR MiaPaCa2 cells (Figure S3A). No change was observed with GEMR in either cell line for radiolabelled palmitic acid uptake (Figure S3B).

### GEMR increases stearoyl-CoA desaturase 1 (SCD1)-driven desaturation in Panc1 cells

SCD1 is a Δ9-desaturase that inserts a carbon-carbon double bond at the 9^th^ carbon from the carboxylate end of either palmitic (16:0) or stearic (18:0) acid to form palmitoleic (FA 16:1*n*-7, *cis*) or oleic acid (FA 18:1*n*-9, *cis*) respectively (where the *n-x* refers to the location of the carbon*-*carbon double bond from the methyl end). Immunoblotting revealed increased abundance of SCD1 in Panc1 GEMR cells (Figure 2C). This increase in SCD1 protein abundance could explain increased levels of MUFA-containing lipids in Panc1 GEMR cells; however, recent work has demonstrated the presence of FADS2-derived isomeric MUFA in other cancer types ^35,36^ that contain carbon*-*carbon double bonds at locations distinct from SCD1-derived MUFA (Figure 2A, red/orange labels). Such isomers cannot be differentiated using the LC-MS/MS method used for untargeted lipidomics analysis in Figure 1A&B. To measure SCD1-derived fatty acids specifically in our GEMR PDAC cell lines we performed ozone-enabled fatty acid discovery (OzFAD) ^37^ on AMP^+^-derivatised MUFA (Figure 2E). These data showed a statistically significant increase in SCD1-derived FA 16:1*n*-7 in Panc1 GEMR whereas SCD1-derived FA 18:1*n*-9 was increased in MiaPaCa2 GEMR cells. To confirm these findings, SCD1 activity was measured in both cell lines by measuring desaturation of uniformly labelled ^13^C_16_ palmitate into ^13^C_16_-FA 16:1*n*-7 after 72 h. Baseline levels of FA 16:0 were not different between CON and GEMR cells of either cell line (Figure S4A&B). Despite no change in labelled palmitate uptake in Panc1 cells (Figure S4C), increased FA 16:1*n-*7/FA 16:0 was detected in GEMR Panc1 cells indicating enhanced SCD1 activity (Figure 2F). In contrast, no difference was observed in SCD1 activity for GEMR MiaPaCa2 cells despite a slight increase in labelled palmitate uptake (Figure S4D). Taken together, these data suggest that GEMR upregulates SCD1 abundance and activity specifically in Panc1 cells.

### FADS2-derived monosaturated fatty acids are present in Panc1 cells

Using OzFAD we were also able to detect the presence of Δ6 FADS2-derived *n-*10 isomers of FA 16:1 and FA 18:1 (*c.f.* Figure 2A) in both cell lines, with a greater amount of *n-*10 fatty acids found in Panc1 cells relative to MiaPaCa2 (Figure 2E). Both *n*-10 fatty acid isomers were lower in Panc1 GEMR cells when compared with CON, with no changes detected in MiaPaCa2 GEMR cells. Immunoblotting for FADS2 revealed no difference in abundance between Panc1 CON and GEMR cells; however, FADS2 protein was virtually undetectable in MiaPaCa2 cells (Figure 2G). Only trace amounts of *n-*12 isomers (i.e. 18:0 FADS2-derived MUFA, *c.f.* Figure 2A) were detected in the Panc1 cell lines (data not shown), suggesting a specific upregulation of Δ6 desaturation of FA 16:0 by FADS2. To further elucidate the effect of GEMR on FADS2 we measured its steady-state activity by tracing uniformly labelled ^13^C palmitate into 16:1*n-*10 in both cell lines. No change in FADS2 activity was observed between CON and GEMR Panc1 cells and a small but statistically significant decrease in FADS2 activity was seen in MiaPaCa2 GEMR cells (Figure 2H). Overall, these data suggest that there is no specific increase in FADS2 abundance or *n-*10 MUFA synthesis activity with GEMR in either cell line.

### SCD1 and FADS2 compensate for each other when the alternate is knocked down in Panc1 cells

To investigate if targeting lipid synthesis can improve the response of PDAC cells to GEMC treatment we performed knockdown of key enzymes using small interfering RNA (siRNA). We targeted the genes of key enzymes in lipid synthesis pathways including *ACACA* (*i.e.*, the gene encoding ACC1), *FASN* (FAS), *SCD* (SCD1), and *FADS2*. We tested single siRNA targets against *ACACA* and *FASN* and confirmed knockdown in Panc1 & MiaPaCa2 cells (Figure 3A). Knockdown of either *ACACA* or *FASN* also resulted in large decreases in *de novo* lipogenesis as measured by tracing radiolabelled glucose into the lipid fraction of CON Panc1 & MiaPaCa2 cell lines (Figure 3B). Two different siRNA targets were tested for *SCD* and *FADS2*, with either showing good efficacy in the knockdown of their target (Figure 3C). Therefore, both siRNAs for *SCD* and *FADS2* were used in combination throughout.

**Figure 3:**
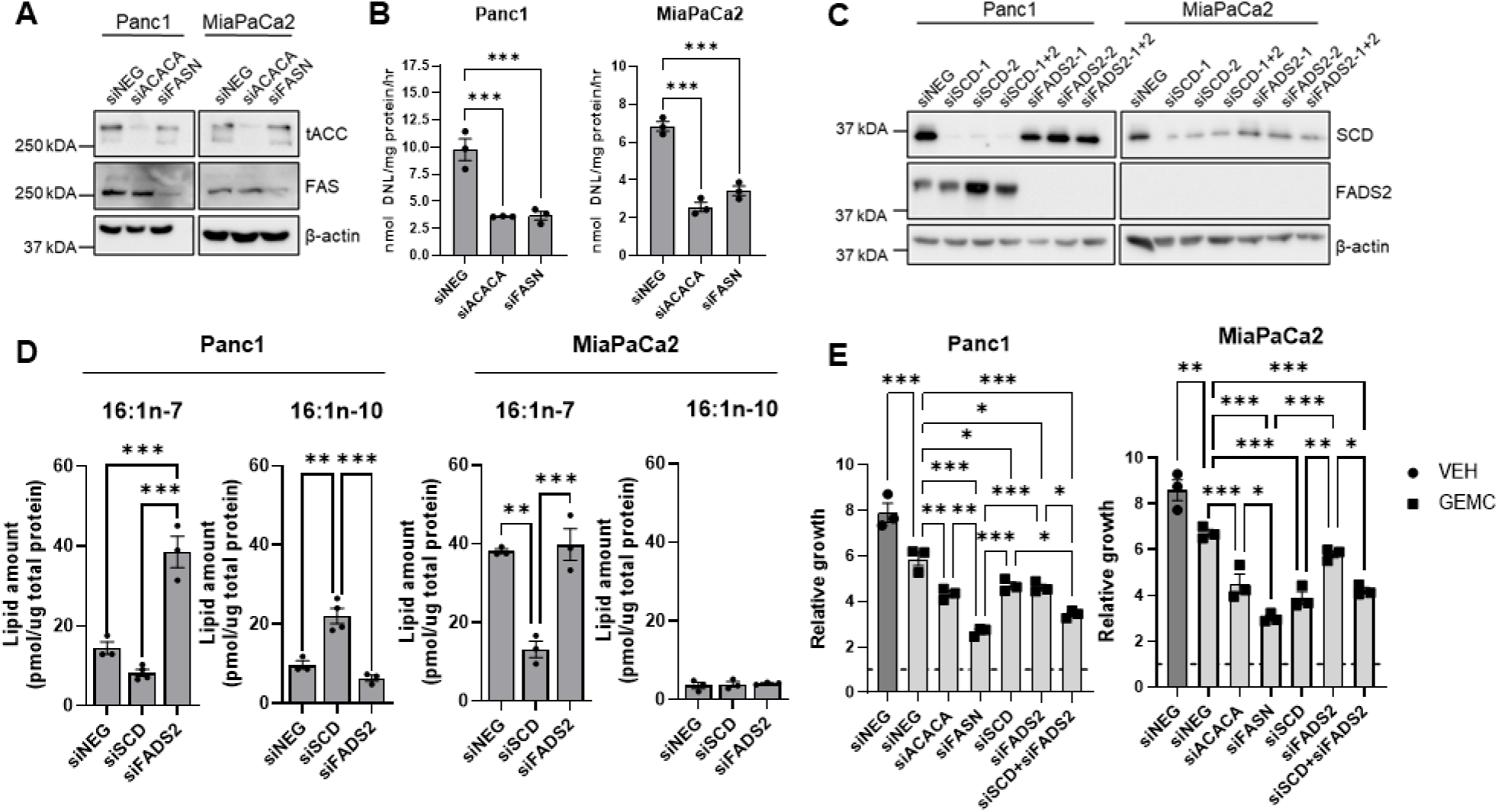
A. Immunoblotting for ACC & FAS and **B** measurement of radiolabelled glucose into lipid fraction following knockdown of *ACACA* or *FASN*. **C** Immunoblotting for SCD1 & FADS2 or **D** quantification of 16:1 isomers measured by liquid chromatography-mass spectrometry following knockdown of *SCD* or *FADS2*. All measurements were performed following 72 h of siRNA transfection. **E** effect of 48 h GEMC treatment (Panc1 1 µM, MiaPaCa2 100 nM, or vehicle, VEH) on relative cell growth following 72 h knockdown of lipid synthesis enzymes. Cell growth was measured by crystal violet assay and normalised to start of siRNA knockdown (dashed line). Statistical significance was determined by **B** unpaired t test, or **D**&**E** one-way ANOVA with Tukey posthoc correction, \**P* < 0.05 ** *P* < 0.01. *** *P* < 0.001

We next measured the impact of SCD1 and FADS2 knockdown on the amount of 16:1 isomers by LC-MS/MS. In MiaPaCa2 cells, SCD1 knockdown had the intended effect where it significantly decreased FA 16:1*n-*7 levels (Figure 3D). Knockdown of FADS2 in MiaPaCa2 cells did not affect the amount of FA 16:1*n-*10; however, levels of FA 16:1*n-*10 (Figure 2E) and the abundance of FADS2 protein (Figure 2G&3C) are very low in this cell line. In contrast, knockdown of either desaturase in Panc1 cells did not decrease the expected isomeric MUFA product; instead, the alternate desaturation product increased in abundance (Figure 3D). For example, SCD1 knockdown in Panc1 cells resulted in a large increase in FA 16:1*n-*10, and while the amount of FA 16:1*n-*7 content did decrease relative to si*NEG* with SCD1 knockdown this reduction was not statistically significant. Likewise, the knockdown of FADS2 resulted in a large increase in FA 16:1*n-*7. Together these data suggest that each desaturase can compensate when the alternate is inhibited in Panc1 cells.

### Knockdown of lipogenic enzymes improves GEMC efficacy in PDAC cells

Post validation of knockdown targets, we assessed if reducing the activity of the different lipid enzymes could improve GEMC chemotherapy efficacy on cell growth in Panc1 & MiaPaCa2 cells (Figure 3E). Target enzymes were knocked down in Panc1 & MiaPaCa2 cells for 72 h prior to 48 h of treatment with vehicle (VEH) or a dose of GEMC tailored to induce a statistically significant decrease in si*NEG* cell growth (Panc1 1 µM, MiaPaCa2 100 nM). Knockdown of *ACACA*, *FASN*, *SCD* or *FADS2* combined with GEMC treatment resulted in decreased cell growth in Panc1 cells when compared with GEMC-treated si*NEG* controls, with si*FASN* producing the greatest effect (Figure 3E). *ACC, SCD* or *FADS2* knockdown each had an equivalent impact on cell growth. However, co-targeting desaturases using si*SCD* and si*FADS2* produced a significantly greater decrease in Panc1 cell growth than targeting either enzyme alone, with the effect of dual desaturase knockdown on cell growth being comparable to that of si*FASN* (Figure 3E). All reported decreases in cell growth following concomitant knockdown and GEMC treatment were above that for knockdown alone for all targeted genes except *FASN*, which had a similar impact on growth with or without GEMC treatment (Figure S5A). Knockdown of key lipid synthesis enzymes in combination with GEMC treatment in MiaPaCa2 cells showed a similar trend to that of Panc1, except that si*FASN* or si*SCD* produced a comparable decrease in relative cell growth a (Figure 3E). Knockdown of *FADS2* did not significantly decrease cell growth in MiaPaCa2 cells, and, not surprisingly, co-targeting both desaturases with si*SCD* and si*FADS2* in MiaPaCa2 cells produced no additional effect when compared with si*SCD* (Figure 3E). The effect of combined knockdown with GEMC treatment was above that of knockdown only for *ACACA* and *FADS2* in MiaPaCa2 cells (Figure S5B). We also measured the effect of knockdown of lipid synthesis enzymes in GEMR cells; however, only si*FASN* resulted in decreased cell growth on either cell line (Figure S5C&D). Collectively, these data suggest that targeting key enzymes involved in fatty acid synthesis can successfully sensitise GEMC-naïve PDAC cells to GEMC treatment, and that where alternate desaturase activity exists co-targeting both desaturases is necessary to have the greatest impact on cell growth.

### Combination GEMC/paclitaxel resistance also upregulates lipid synthesis

GEMC monotherapy has long been the first-line treatment for metastatic PDAC, but is now more typically given in combination with nab-paclitaxel ^38^. To test the effect of combination chemotherapy on *de novo* lipogenesis in PDAC we attenuated Panc1 and MiaPaCa2 cells to grow in GEMC/paclitaxel (PTX) by a similar regime to that of GEMR induction using a ratio of 10:1 (GEMC:PTX) to approximate the typical ratio given to PDAC patients ^39,40^. In Panc1 cells the final concentration was 28 nM GEMC (2.8 nM PTX), and 8 nM GEMC (0.8 nM PTX) for MiaPaCa2 cells. Combination therapy attenuated (CombAT) Panc1 and MiaPaCa2 cells showed improved growth and proliferation in media containing 15 nM GEMC/1.5 nM PTX when compared to naïve control cells, suggesting that resistance was present in both cell lines (Figure S6). We next performed untargeted lipidomics to determine relative changes in lipid species across both cell lines (Figure 4A&B). These data revealed extensive increases across 166 lipid species in Panc1 CombAT cell lines compared with CON, including increases in many triacylglycerol (TG) species and total TG (Figure 4A). Increases were also seen in several TG species in CombAT MiaPaCa2 cells, with 30 lipid species increased in total when compared with CON (Figure 4B). Together these data suggest that the addition of PTX to GEMC chronic treatment does not abrogate the lipogenic phenotype observed with monotherapy GEMR, and, in fact, may enhance *de novo* lipogenesis in both cell lines.

**Figure 4:**
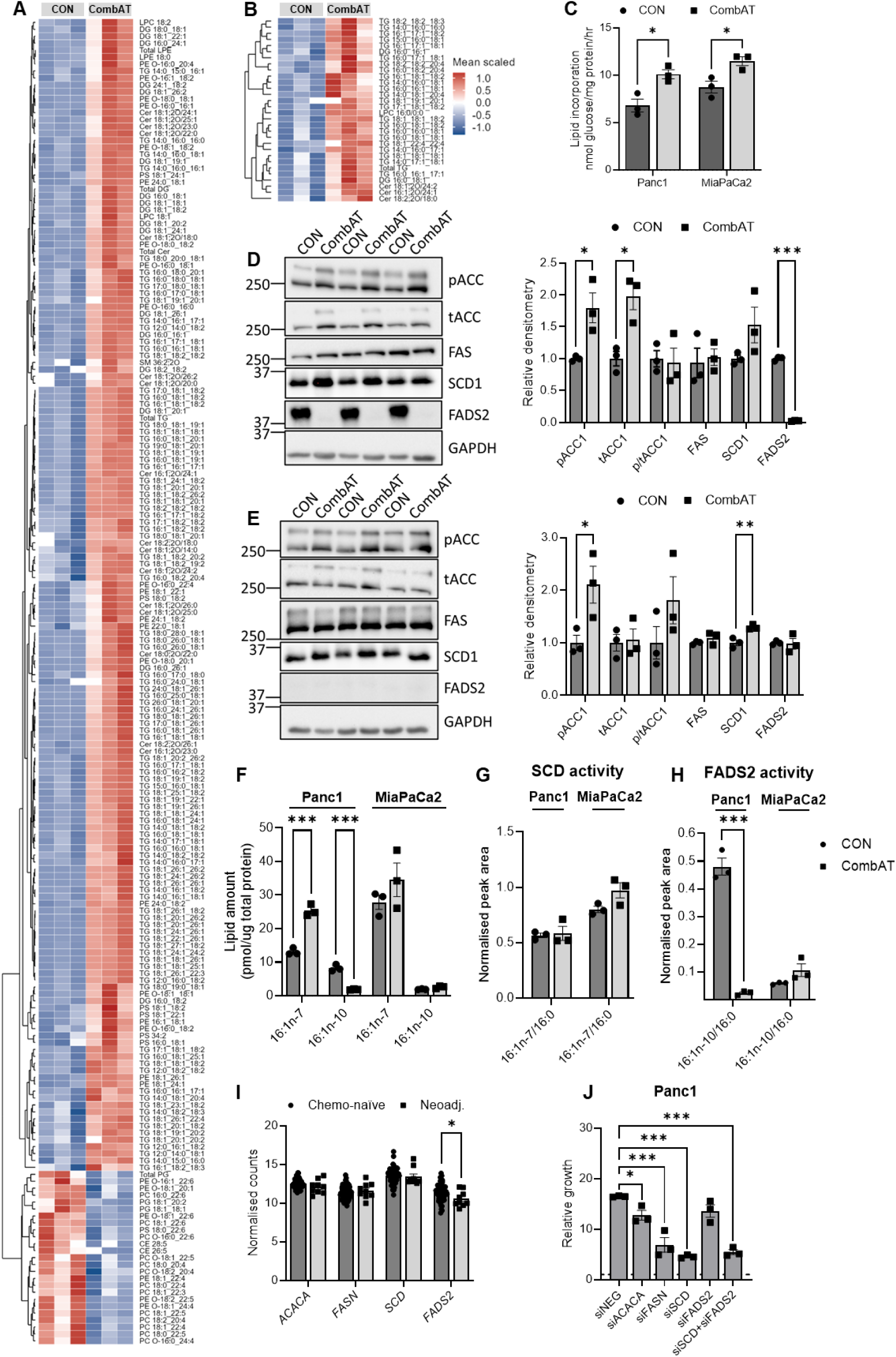
Untargeted lipidomics of both **A** Panc1 and **B** MiaPaCa2 control (CON) and combination therapy attenuated (CombAT) cell lines (unpaired t-test, *P* < 0.01). **C** *De novo* lipogenesis measured by the incorporation of radiolabelled glucose into the lipid fraction. Immunoblotting for total and activated key lipid synthesis enzymes in **D** Panc1 and **E** MiaPaC2 CON and CombAT cells. **F** 16:1 fatty acid isomers measured by LC-MS of AMP^+^-derivatised fatty acids. Measurement of **G** SCD1 and **H** FADS2 activity in Panc1 and MiaPaCa2 CON and CombAT cell lines by tracing uniformly stable-labelled palmitic acid into isomeric MUFA. **I** Analysis of PDAC tumour RNAseq data published by Zhou and colleagues obtained from patients who underwent surgical resection prior to chemotherapy (chemo-naïve) or received GEMC/nab-paclitaxel neoadjuvant treatment (Neoadj.) before tumour resection. **J** Effect of 72 h knockdown of lipid synthesis enzymes on Panc1 CombAT cell growth. Statistical significance was determined by unpaired t test (**A-I**) or one-way ANOVA with Tukey posthoc comparisons with *si*NEG only (**J**). \**P* < 0.05 ** *P* < 0.01. *** *P* < 0.001.

We next assessed if increased *de novo* lipogenesis persisted in our CombAT cell lines, with both Panc1 and MiaPaCa2 CombAT cells demonstrating increased radiolabelled glucose incorporation into the lipid fraction (Figure 4C). Immunoblotting for key lipogenic enzymes showed increases in total and phosphorylated ACC1 in Panc1 CombAT cells (Figure 4D), alongside increased levels of phosphorylated ACC1 in MiaPaCa2 combat cells (Figure 4E). A small but statistically significant increase in SCD1 abundance was also detected in MiaPaCa2 CombAT cells.

### Combination chemotherapy resistance decreases FADS2 abundance and activity in Panc1 cells

Surprisingly, FADS2 protein abundance was substantially decreased in Panc1 CombAT cells (Figure 4D). Similar to GEMR cell lines, little to no FADS2 protein was detected in MiaPaCa2 CombAT or CON cells (Figure 4E). To confirm decreased FADS2 abundance in Panc1 CombAT cells we measured FA 16:1 isomers in both cell lines by LC-MS/MS and found increased FA 16:1*n-*7 in Panc1 CombAT cells alongside substantially decreased FA 16:1*n-*10 (Figure 4F). Also observed in Panc1 CombAT were changes in PUFA profile consistent with FADS2 downregulation, including decreases in FA 22:6*n-*3, FA 18:3*n-*6, FA 20:4*n-*6, FA 22:4*n-*6 and FA 22:5*n-*6 when compared with CON (Figure S7). No changes were detected in FA 16:1 isomeric profile in MiaPaCa2 CombAT cells when compared to CON (Figure 5F), alongside no changes in PUFA levels (Figure S7C). To further confirm these findings, we measured steady-state desaturase activity by tracing uniformly labelled ^13^C-palmitate into both cell lines over 72 h. Neither cell line exhibited any differences between CON and CombAT cells in unlabelled FA 16:0 content (Figure S8A&B). However, Panc1 CombAT cells had more labelled palmitic acid (relative to unlabelled palmitic acid) while MiaPaCa2 CombAT cells had less (Figure S8C&D respectively). No differences were seen for SCD1 activity as measured by the ratio of labelled FA16:1*n-*7 to FA 16:0 in Panc1 or MiaPaCa2 CombAT cells (Figure 4G). A substantial decrease in FA 16:1*n*-10/16:0 was detected in Panc1 CombAT cells, indicating a reduction in FADS2 activity, with no change in FA 16:1*n-*10/FA 16:0 observed for MiaPaca2 CombAT cells (Figure 4H). Also apparent in Panc1 CombAT cells was a marked decrease in FAO compared with CON cells (Figure S9). Taken together, these data indicate that chronic combination GEMC/PTX treatment strongly downregulates FADS2 abundance and activity in the Panc1 cell line. To explore if combination GEMC/PTX therapy might exhibit the same effect on lipid metabolism in human PDAC patients, we analysed recently published RNA-seq data obtained from PDAC patients who received no treatment (chemo-naïve) or GEMC/nab-paclitaxel neoadjuvant chemotherapy prior to tumour resection ^41^. The data showed no difference in the expression of *ACACA, FASN or SCD* between chemo-naïve or neoadjuvant-treated tumours; however, the neoadjuvant group demonstrated a small but statistically significant decrease in *FADS2* expression when compared with chemo-naïve tumours (Figure 4I).

Given the effect of combination therapy resistance on FADS2 abundance and fatty acid isomer profile we sought to understand if co-targeting both desaturases was still necessary to decrease in cell growth in Panc1 CombAT cell lines. Knockdown of *ACACA*, *FASN*, or *SCD* for 72 h in Panc1 CombAT cells resulted in a statistically significant decrease in cell growth when compared to negative siRNA control (si*NEG*), with si*FASN* or si*SCD* producing the largest decreases in cell growth (Figure 5J). Panc1 CombAT cells were insensitive to *si*FADS2 knockdown, and no additional effect on cell growth was observed with co-knockdown of SCD. These data suggest that therapeutically targeting SCD1 in combination with GEMC/PTX chemotherapy could be a viable strategy for enhancing cytotoxicity in PDAC.

## Discussion

We sought to characterise metabolic adaptations to chemotherapy exposure in PDAC and found enhanced *de novo* lipogenesis in PDAC cells with resistance to both monotherapy GEMC and combination GEMC/PTX chemotherapy. Enhanced *de novo* lipogenesis is a known feature of many cancers, including pancreatic cancer ^18^, where increased expression of lipogenic genes is correlated with poorer survival and greater resistance to chemotherapy ^17^. Herein we provide the first report of enhanced *de novo* lipogenesis specifically in GEMC resistance in PDAC, providing a novel target for the disease that could potentially improve treatment efficacy. We also provide the first report of FADS2-derived *n*-10 MUFA in PDAC (Figure 2E). The ability of cancer cells to exhibit diversity in isomeric lipids has recently been described in other cancer cell types including liver & lung cancer cell lines more extensively in prostate cancer ^35,36^. This metabolic and isomeric diversity is proposed to promote metabolic plasticity enabling cancer cells to survive and proliferate in challenging tumour microenvironments depleted in bioenergetic fuels and oxygen.

In this work we specifically found upregulation of ACC1 (through increased total abundance or decreased phosphorylation at Serine-79), FAS and SCD with GEMR. FAS is commonly expressed at high levels in a variety of cancers of human tissues ^23–25^ and *FASN* overexpression correlates to poorer survival and increased GEMC resistance in PDAC ^17^. High FAS levels have also been found in the serum of patients with PDAC or intraductal papillary mucinous neoplasms ^42^. We found that genetically inhibiting FAS (Panc1 & MiaPaCa2), SCD1 (MiaPaCa2) or SCD1 & FADS2 (Panc1) in combination with GEMC produced the greatest impact on PDAC cell growth (Figure 4E). Previous work demonstrated that inhibition of FAS by Orlistat induces ER stress and acts synergistically with GEMC to decrease PDAC cell growth and reduce tumour burden in xenograft models of PDAC ^17^. Orlistat has also been reported to attenuate oxaliplatin resistance in colorectal cancer ^43^. This synergistic effect of Orlistat with chemotherapy *in vivo* may be related to a direct effect on FAS but may also arise from the dual action of Orlistat on lipases that further inhibit the availability of free fatty acids for use by cancer cells. However, Orlistat has poor bioavailability, is contraindicated in pancreatic disease, and can cause pancreatitis in susceptible patients ^44^, all of which potentially limit its therapeutic use in PDAC. Other small molecule inhibitors of FAS have had some success in decreasing cancer cell growth, including in breast, colon, lung, colorectal, bladder, ovarian, gastric, endometrial, prostate, and gastrointestinal cancers as well as in melanoma and lymphoma (reviewed extensively by ^23^). Cerulenin has previously been shown to re-sensitise cisplatin*-*resistant ovarian cancer cell lines to treatment ^45^, while its more stable derivative C75 sensitises breast cancer cells to cisplatin treatment ^46^. However, both cerulenin and C75 exhibit significant off-target and side effects that preclude their clinical use ^23^. Newer FAS inhibitors TVB-3166 and fasnall show promise for synergistically inhibiting cell viability when combined with tubulin*-*stabilising drugs in taxane-resistant prostate cancer ^47^. Mouse and human*-* derived mutant *KRAS^G12D^* lung cancer models are sensitive to genetic or TVB-3664 inhibition of FAS ^48^. Moreover, high *FASN* and *SCD* expression are correlated with mutant *KRAS^G12D^* in lung cancer ^48^, which could potentially explain the differences between the two PDAC cell lines examined herein – Panc1 cells similarly contain *KRAS^G12D^* and GEMR increased expression and activity of both lipogenic enzymes, while MiaPaCa2 cells contain *KRAS^G12C^* (Table S1). In any event, targeting FAS could provide therapeutic benefits to PDAC patients when combined with GEMC-based therapy; however, a small molecule inhibitor with good efficacy and a low side effect profile would need to be developed.

SCD1 has been described as an oncogene in lung cancer ^49^ and explored as a potential therapeutic target for several cancers including PDAC (reviewed by ^50^). SCD1 is highly expressed in the exocrine pancreas ^51^ and increased expression of SCD1 in cancer-associated fibroblasts correlates with decreased PDAC survival time and GEMC resistance ^16^. Higher levels of FA 16:1 and 18:1 are associated with aggressive subtypes of PDAC immortalised cells; however, positions of unsaturation (*i.e.*, fatty acid isomers) were not determined in this work ^31^. Inhibition of SCD1 using the small molecule inhibitor A939572 induces the unfolded protein response and decreases the growth of pancreatic tumours in KPC mice and in EPCAM^+^ organoids derived from KPC mouse tumours ^52^. In this work, we also discovered significant FADS2-derived *n*-10 MUFA desaturase products in the Panc1 cell line (Figure 2E). There is limited exploration of FADS2 as a therapeutic target in cancer and existing data has largely focused on its role in Δ6 desaturation of PUFA ^53,54^. FADS2 inhibition by small molecule inhibitor sc-26196 suppresses prostaglandin E2 production via arachidonic acid and sensitises glioblastoma cells to radiotherapy in a xenograft model ^55^. Genetic or pharmacological FADS2 inhibition also suppresses colon, lung, melanoma, and cholangiocarcinoma cancer cell tumour growth in xenograft and genetically engineered models ^56–58^. Pharmacologic or genetic dual-inhibition of SCD1 and FADS2 decreases tumour and metastatic lesion growth and cancer stem cell formation in ovarian cancer while reducing resistance to platinum-based chemotherapy ^54^. Our work and others ^35,36^ observed overcompensation by the alternative desaturation pathway when SCD1 or FADS2 are individually inhibited in cancer cells (Figure 4D). Indeed, this same compensatory desaturase effect might explain the observed decrease in 16:1*n*-10 and 18:1*n*-10 in Panc1 GEMR cells, where SCD1 abundance, activity, and MUFA products were all increased when compared with Panc1 CON cells (Figure 2C, E, &F). Similarly, an increase in *FADS2* expression has been observed when cell lines resistant to SCD1-inhibitors (A549 & HeLa) are treated with A939572 ^59^. We show suboptimal inhibition of PDAC cell growth when only one desaturase is silenced alongside co-treatment with GEMC (Figure 4E). Therefore, the use of SCD1 inhibitors might be suboptimal in the presence of significant FADS2 abundance or activity; however, the extent of FADS2 protein abundance or activity in PDAC tumours on a population level is not yet fully known.

This phenomenon of competing inhibitory pathways has previously been described and is thought to enable enhanced plasticity at the expense of maximised proliferation ^36,60,61^. Such competing pathways might also promote cancer cell survival in PDAC as the hypoxic, nutrient-depleted tumour microenvironment is particularly enhanced by the dense fibrosis associated with the disease ^62^. Desaturation is a highly energy-demanding process, and desaturases are NADPH, O_2_, and cytochrome b5 dependent ^63^ - all of which are likely in short supply in the tumour microenvironment. Specifically in PDAC, mutant *KRAS* regulates GOT1 synthesis of NADPH from glutamine to maintain redox status ^64^, but feasibly this pathway could also support high NADPH levels to maintain enhanced lipid synthesis and desaturation. Though highly energy-demanding, desaturation would allow the sequestration of highly lipotoxic SFA such as FA 16:0 and FA 18:0 into less toxic MUFA in lipogenic cells, thus preventing cell death ^65^. Increased desaturation of lipids also modulates cell membrane fluidity and is highly associated with epithelial-to-mesenchymal transition (EMT). This process is crucial for cell migration and metastatic initiation ^66,67^, and FADS2-derived FA 16:1*n-*10 specifically promotes cell migration in melanoma cells ^68^. Expression of EMT transcription factor Zeb1 has been shown to promote increased PUFA to MUFA ratio within cancer cells, with downregulation of Zeb1 suppressing FADS2 and enhancing SCD1 & FAS expression ^69^. MUFA are also less susceptible to peroxidation by reactive oxygen species (ROS) than PUFA, and given that ROS levels are often high in PDAC and other cancers ^70,71^ increased MUFA would limit both ROS and cell death from iron*-*driven lipid peroxidation (*i.e.* ferroptosis). Indeed, cells cultured with exogenous sources of MUFA have suppressed ferroptosis ^72^ and SCD1 overexpression protects ovarian cancer cells from ferroptosis-mediated cell death ^73^. Similarly, pharmacological inhibition of SCD1 sensitises cancer cells expressing high levels of Zeb1 to ferroptosis ^69^. In short, despite the high energy demands associated with lipid desaturation the process likely provides significant benefits for cancer cell survival that outweigh the energetic cost.

Our finding that combination GEMC/PTX resistance strongly and very specifically downregulates FADS2 protein abundance and activity in Panc1 cells was unexpected. Single-dose combination GEMC/nab-paclitaxel downregulates *de novo* lipogenesis in PDAC cells via the AMP-activated protein kinase/ sterol regulatory element-binding protein 1 (SREBP1) axis ^74^. However, PTX treatment has also previously been shown to increase FADS2 protein abundance in lung adenocarcinoma cells ^75^. The mechanism by which PTX combined with GEMC specifically downregulates FADS2 protein levels and activity is unknown, and FADS2 shares similar transcriptional factors to ACC, FAS and SCD1 ^76^. Upregulated *FADS2* expression and increased sapienic acid biosynthesis can be driven in multiple cancer cell lines through increased mammalian target of rapamycin (mTOR) signalling and SREBP ^77^; however, little else is known about transcriptional or post-translational regulation of FADS2 abundance and activity. Nevertheless, our results strongly suggest that combining SCD1 inhibition with GEMC/PTX chemotherapy could be an effective broad-spectrum strategy for decreasing PDAC cell growth that could benefit patients both with or lacking significant dual desaturase activity. This could be important for accelerated translational efforts as a therapeutic target as only a single small molecule inhibitor is currently available for FADS2 (sc-26196), and its *in vivo* safety and efficacy profile is yet to be established in humans. Importantly PDAC patients who received GEMC/PTX neoadjuvant treatment show decreased *FADS2* expression (Figure 5I), further highlighting the potential translatability of our findings.

In summary, we have identified that *de novo* lipogenesis is increased in GEMC-resistant PDAC cells including the presence of an alternate desaturation pathway in Panc1 cells. We identify several key lipid metabolic pathways including ACC1, FAS, SCD1 and FADS2 that could be targeted to improve GEMC efficacy on cell growth. Importantly, we have determined that SCD1 and FADS2 can compensate for each other in PDAC, and that where dual desaturase activity exists both desaturases must be targeted to provide the best effect. Finally, we provide evidence that combination GEMC/PTX therapy strongly and specifically downregulates FADS2 expression and activity and increases the efficacy of SCD1 inhibition in PDAC cells with dual desaturase activity. Therefore, we conclude that co-targeting lipid synthesis in PDAC could be a viable strategy for improving the efficacy of both GEMC monotherapy and combination GEMC/PTX therapy in patients.

## Materials and method

### Materials

Gemcitabine hydrochloride, paclitaxel and foetal calf serum (FCS) were purchased from BioStrategy (Melbourne, VIC, Australia). High glucose Dulbecco’s Modified Eagle’s medium (DMEM) was purchased from ThermoFisher Scientific (Melbourne, VIC, Australia). All solvents and additives used for liquid chromatography-mass spectrometry (LC-MS) were HPLC grade or higher and were purchased from ThermoFisher Scientific. Crystal violet and LC-MS grade water were purchased from Merck Life Sciences (Melbourne, VIC, Australia). All lipid internal standards were purchased from Merck Life Sciences except diacylglycerol 15:0/15:0 (Nu-Chek Prep, Elysian, MN, USA) and d5-triacylglycerol 16:0/16:0/16:0 (C/D/N Isotopes, Quebec, Canada). Antibodies for immunoblotting were purchased from Cell Signaling Technology (Danvers, MA, USA), Abclonal (Woburn, MA, USA) Abcam (Melbourne, VIC, Australia), or Santa Cruz Biotechnology (Dallas, TX, USA) as indicated in Table S1. MUFA isomer standards were purchased from Sapphire Bioscience (Sydney, NSW, Australia).

### Cell culture & chemotherapy resistance generation

Panc1 and MiaPaCa2 cells from ATCC were cultured in high glucose DMEM containing 4.5 g/L glucose and were supplemented with 10% FCS. Cells were cultured under 5% CO_2_ at 37°C and routinely passaged every 2-3 days (when ∼80-90% confluent). GEMC resistance was induced in PDAC cell lines through increasing doses over 10-12 weeks. Each cell line was treated with the highest tolerable dose of gemcitabine for 48 h and allowed to recover until normal proliferation was observed (1-3 weeks). This process was repeated a minimum of three times before comparing the half-maximal inhibitory concentration (IC_50_) with untreated cells of the same PDAC line. If the IC_50_ in the GEMC-treated group was higher than CON then resistance was determined to have been established. If not, then the concentration of GEMC was increased, and the process repeated. CombAT cells were generated similarly through repeat escalating doses of GEMC/PTX (10:1 v/v) for 12 weeks.

### Measurement of half-maximal inhibitory concentration

The IC_50_ was determined using a crystal violet assay. Cells were seeded at 1×10^4^ cells/well into a 96-well plate in triplicate and allowed to adhere overnight. The next day cells were treated with serial dilutions of GEMC (phosphate buffered saline, PBS, as vehicle) or GEMC:PTX (10:1, with PBS+0.1% ethanol as the vehicle). The IC_50_ was measured after 48 h by crystal violet assay. Media was removed from adherent cells and cells were washed once with PBS. Cells were fixed in a solution of 0.5% w/v crystal violet in 50% v/v methanol for ≥10 min. After fixation, cells were washed with water to remove excess crystal violet and allowed to dry overnight. Stained adherent cells were then solubilised in 1% w/v SDS by vigorous shaking for several hours and the absorbance was read at 570 nm. Absorbance values were normalised to the vehicle and the IC_50_ was calculated by a non*-*linear regression by fitting the log_10_ transformed drug concentration against relative cell viability. Statistical significance was determined by comparing a four-parameter curve fit using the extra sum of squares F test (*P* < 0.05).

### BODIPY staining and confocal imaging

For BODIPY staining and confocal imaging, cells were seeded in a 96 well plate at 10,000 cells/well and allowed to grow to near-confluency. Cells were then fixed with 4% paraformaldehyde for 10 min at room temperature, followed by PBS washes. Permeabilization was performed for 10 min using a saponin*-*based permeabilization/blocking buffer (0.2% (w/v) saponin, 4% (v/v) FCS, 1% (w/v) bovine serum albumin (BSA), and 0.02% (v/v) sodium azide in PBS). Next, cells were incubated for 2 h with Alexa Fluor® 647 anti-Tubulin*-*α antibody (1:250 dilution, clone 10D8, BioLegend) prepared in the permeabilization/blocking buffer. After several washes with PBS, 200 nM BODIPY 493/503 (Cat. No. 25892, Cayman Chemical) and Hoechst 33342 (10 μg/mL, B2261-25MG, Sigma-Aldrich) were added in permeabilization buffer for 10 min at room temperature, followed by PBS washes. Confocal images were acquired using an LSM900 confocal laser scanning microscope with a 40x objective. Lipid droplets per cell and droplet size were quantified using Fiji software.

### Lipid extraction

Lipids were extracted from cells grown to 80-90% confluency in 6 well plates using a modified methyl *tert*-butyl ether (MTBE) method ^78,79^. Adherent cells were washed with ice-cold PBS and scraped into methanol containing 0.01% butylated hydroxytoluene and 100 pmol of internal standards (phosphatidylcholine 19:0/19:0, phosphatidylethanolamine 17:0/17:0, phosphatidylserine 17:0/17:0, lysophosphatidylcholine 17:0, lysophosphatidylethanolamine 14:0, phosphatidylglycerol 17:0/17:0, phosphatidic acid 17:0/17:0, ceramide d18:1/17:0, dihydrosphingomyelin 12:0, cholesteryl ester 22:1, diacylglycerol 17:0/17:0, D_5_-triacylglycerol 16:0/16:0/16:0 and cardiolipin 14:0/14:0/14:0/14:0). An empty well was also scraped for use in background subtraction. To this, MTBE was added at a ratio of 3:1 v/v, and extracts were rotated overnight at 4°C. Following overnight rotation, 1 part of ice-cold 150 mM ammonium acetate was added, samples were vortexed vigorously and centrifuged at 2000 *x g* for 5 min. The upper organic phase containing lipids was transferred into a 1.5 mL autosampler vial and dried under N_2_ at 37°C Dried lipids were reconstituted in chloroform:methanol:water (60:30:4.5 v/v/v), transferred to a sleeved vial, and stored at −20°C until analysis. The aqueous phase containing the protein pellet was dried under N_2_ and digested with 1M NaOH overnight at 4°C. Digested protein was diluted 1:2 v/v with water and protein concentration was determined using a Pierce BCA assay kit (ThermoFisher Scientific)

### Untargeted lipidomics

Untargeted lipidomics was performed as previously described ^80^ using a Dionex Ultimate 3000 LC pump and Q Exactive Plus mass spectrometer equipped with a heated electrospray ionization (HESI) source (ThermoFisher Scientific, Waltham, MA, USA) ^81–83^. Lipids were separated on a core-shell hybrid C18 reverse-phase column (ACQUITY C18 CSH, 2.1 × 100 mm, 1.7μm pore size, Waters Corp., Milford, MA) using a binary gradient, where mobile phase A consisted of acetonitrile:water (6:4 v/v) and B of isopropanol: acetonitrile (9:1 v/v). Both mobile phases A and B contained 10 mM ammonium formate and 0.1% formic acid. The autosampler was kept chilled at 10°C, the column was heated to 60°C, and the flow rate was maintained at 0.26 ml/min. Source conditions were as follows: a spray voltage 4.0 kV in positive ion mode and 3.5 kV in negative ion mode, capillary temperature 290°C, S lens RF 50, and auxiliary gas heater temperature of 250°C. Nitrogen was used as both source and collision gas, with the sheath gas flow rate set at 20 and the auxiliary gas flow rate at 5 (arbitrary units). Data were acquired in full scan/data-dependent MS2 mode (full-scan resolution 70,000 FWHM, max ion injection time 50 ms, scan range m/z 200–1,500), with the 10 most abundant ions being subjected to collision*-*induced dissociation using an isolation window of 1.5 Da and a normalized stepped collision energy of 15/27 eV. Product ions were detected at a resolution of 17,500. An exclusion list for background ions was developed using extraction blanks, and mass calibration was performed in both positive and negative ionization modes before analysis to ensure mass accuracy of 5 ppm in full-scan mode.

Lipids were identified using MS-DIAL v4.9 ^84^. Lipids were detected in both positive and negative ionization modes using a minimum peak height of 1×10^4^ cps, an MS1 tolerance of 5 ppm and MS2 tolerance of 10 ppm, and a minimum identification score of 50%. Identified peaks were aligned with a retention time (RT) tolerance of 0.5 min. Exported aligned data were background subtracted, quantified from internal standards, and normalised to protein using the statistical package R.

### *De novo* lipogenesis assay using [U-14]C labelled glucose

Cells were seeded in 6 well plates and allowed to grow to near-confluency (80-90%, ∼1-2×10^6^ cells/well). Briefly, cells were washed with PBS and incubated in DMEM containing 1 g/L D-glucose and 2 μCi/mL [U-14C]C-glucose (Perkin Elmer, Sydney, NSW, Australia) for 2 h at 37°C. Following this, cells were washed and scraped into ice-cold PBS. Cells were lysed by passing repeated through a fine gauge needle, and an aliquot was taken for protein determination by BCA assay. Lipids were extracted from the remaining sample by Folch extraction ^85^. The lower lipid-containing organic phase was removed to a new vial and dried under N_2_ at 37°C before being reconstituted in scintillation fluid (Ultima Gold, Perkin Elmer, Sydney, NSW, Australia). ^14^C content was determined by direct DPM assay using a beta counter (TriCarb liquid scintillation counter, Perkin Elmer) and normalised to total protein.

### Western blotting

For immunoblotting of lipid synthesis enzymes cells were grown to near-confluency on 6 well plates (∼1-2×10^6^ cells/well). Cells were washed with ice-cold PBS and scraped into ice-cold radioimmunoprecipitation assay buffer containing protease and phosphatase inhibitors (10 μg/mL phenylmethylsulfonyl fluoride, 10 μg/mL aprotinin, 10 μg/mL leupeptin, 1 mM Na_3_VO_4_, and 10 mM NaF). Cells were lysed by repeated freeze/thawing on ice using liquid N_2_, before being centrifuged at 16,100 *x g* (4°C) for 5 min. The supernatant was removed to a new vial, from which the protein concentration was determined by BCA assay. Lysates were resuspended in laemelli buffer, and 10-15 ug of protein was separated by SDS polyacrylamide gel electrophoresis (10%). Proteins were transferred onto 0.2 μm polyvinylidene fluoride membrane and protein bands visualised by Ponceau S staining. Destained membranes were blocked in 5% w/v skim milk in Tris-buffered saline containing 0.1% Tween20 (TBST) for 1 h, before being incubated in primary antibody (1:1000) overnight (Table S2). Membranes were washed with TBST before being incubated with an appropriate secondary antibody conjugated to horse radish peroxidase (1:10,000, 5% w/v skim milk in TBST) for 1 h. Membranes were washed in TBST and antibodies were detected by enhanced chemiluminescence (Clarity Western ECL substrate, Bio-Rad Laboratories, Sydney NSW, Australia, or SuperSignal West Femto Maximum Sensitivity Substrate, ThermoFisher Scientific) and imaged using a ChemiDoc imaging system (Bio-Rad Laboratories). Densitometry was performed using Image Lab v6.1 (Bio-Rad Laboratories) and normalised to the mean of the control group.

### Hydrolysis and AMP derivatisation of fatty acids

Lipids were extracted from adherent cells grown to near-confluency in 6 well plates using the procedure described above with the addition of 20 nmol of methyl nonadecanoate (Merck Life Science) as an internal standard. Following MTBE extraction of lipids, an aliquot was taken and hydrolysed in 0.6 M KOH in 75% v/v methanol at 60°C for 30 min. Samples were then cooled to room temperature and neutralised with 25% v/v acetic acid. Water and *n*-hexane (3:2 v/v) were added separately and samples were vortexed vigorously before being centrifuged at 2000 *x g* for 5 min. The upper phase containing hydrolysed fatty acids was removed, and the aqueous phase was washed with a second volume of *n*-hexane and combined. The combined organic phase was then dried under N_2_ at 37°C. Dried hydrolysed fatty acids were derivatised with an AMP+ MaxSpec kit (Cayman Chemical, Ann Arbor, MI, USA) as per manufacturer instructions ^86^. Following this, AMP-derivatised fatty acids were re-extracted using MTBE:water (1:1 v/v), and the upper MTBE phase was dried under N_2_ before being resuspended in methanol. Samples were stored at −20°C until analysis. A 37-component fatty acid methyl ester standard (Merck Life Sciences) containing an additional 3.184 nmol of methyl nonadecanoate was concurrently subjected to the same derivatisation procedure and used as an external quality control for fatty acid identification by LC-MS. Derivatised MUFA standards FA 16:1*n-*9,*cis*, FA 16:1*n-*10,*cis*, FA 18:1*n-*7,*cis*, FA 18:1*n-*10*,cis*, and FA 18:1*n-*12,*cis* were also spiked into this mixture to generate retention times (RT) for MUFA isomer identification by LC-MS.

### OzFAD analysis of AMP-derivatised fatty acids

Ozone-enabled fatty acid discovery analysis was conducted as previously described ^37^. Briefly, 10 μL injections of AMP+ derivatized fatty acids were subjected to reversed-phase chromatography using a Waters ACQUITY Class UPLC (Waters Corporation, Milford, USA) fitted with an ACQUITY C18 reversed-phase column (2.1 × 100 mm i.d., particle size 1.7 μm, Waters Corporation) maintained at 60 °C. The 20 min gradient utilised a binary mobile phase system comprised A: water (0.1% formic acid) and B: acetonitrile (0.1% formic acid). Chromatographic detection was conducted using an ion-mobility enabled quadrupole time of flight (QTOF) mass spectrometer (SYNAPT G2-S*i* HDMS, Waters Corporation) previously modified to allow introduction of high concentration ozone into the ion mobility region of the instrument ^87^. Electrospray ionisation parameters were: capillary +3 kV, sampling cone +40 V, source temperature 120 °C, desolvation temperature 550 °C, cone gas 100 L/h, desolvation gas 900 L/h, and nebulizer gas pressure 6.5 bar. Gas flows to the ion mobility region of the instrument were 2.0, 180 and 90 mL/min, for the trap, helium cell and IMS region, respectively. For the ozonolysis reactions, oxygen (≥ 99.999%, Coregas, NSW, Australia) was converted to ozone at concentrations of 220-240 g/Nm^3^ (∼15-17% O_3_ in O_2_) using a high concentration ozone generator (TG-40, Ozone Solutions, Hull, IA). Peak areas for Criegee and aldehyde ions for each monounsaturated fatty acid isomer were combined and normalised to internal standard and total protein (measured by BCA assay).

### Desaturase activity determination

Cells were seeded in 6 well plates at a density of 3×10^5^ cells/well and allowed to adhere overnight. The following day, cell culture media was changed to FCS-free high glucose DMEM containing 12.5 µM ^13^C_16_-labelled or unlabelled palmitate conjugated to 2% fatty acid-free BSA. Palmitate was prepared in cell culture media from 100 mM ethanolic stocks of palmitic acid and conjugated to fatty acid-free BSA by incubating at 55 °C in media for 2 h. Lipids were extracted and derivatised as described above after 72 h of incubation with labelled or unlabelled palmitate. Lipids were identified and analysed as described below for LC-MS analysis of AMP-derivatised fatty acids, with labelled fatty acids identified from [M+H+16.0542]^+^ precursors with the same RT as their unlabelled counterparts as per Table S3.

### LC-MS analysis of AMP-derivatised fatty acids

AMP-derivatised fatty acids were separated on a core-shell C30 reverse phase column (Accucore 2.6 μm, 2.1 x 150 mm, ThermoFisher Scientific) on an Agilent 1290 Infinity II HPLC pump (Agilent Technologies Inc, Santa Clara, CA, USA) using a modified gradient to that described above for OzFAD analysis. Mobile phase A consisted of water and mobile phase B of acetonitrile (both containing 0.1% formic acid), the flow rate was set at 0.4 mL/min throughout and the column heater at 30°C. The gradient was as follows: 0-3.25 min increase from 50% B to 56% B, 3.25-6.5 min increase to 58% B, 6.5-7.5 min increase to 80% B, 7.5-9.5 min increase to 100% B, hold at 100% B for 1.5 mins. The column was then re-equilibrated at 50% B for 4.5 mins. AMP-derivatised fatty acids were detected as positive ions in QTOF-only mode on an Agilent 6560 IM-MS QTOF mass spectrometer (Agilent Technologies). Source conditions were as follows: capillary voltage of 3.5 kV, nozzle voltage of 1 kV and fragmentor at 360 V. Nitrogen was used as the sheath and collision gas, with drying gas set at 5 L/min and sheath gas at 12 L/min (both gases heated to 300°C) and the nebuliser pressure set at 20 psi. Ions were acquired in Auto MS/MS mode at an MS1 mass range of *m/z* 100-1000 and MS2 mass range of *m/z* 50-600. Spectra were acquired in MS1 mode at 5 spectra/s and 200 ms/spectrum, and in MS2 mode at 10 spectra/s and 100 ms/spectrum. Precursor ions were isolated using a window of 1.3 Th, acquisition time of 50 ms/spectra, and collision energy of 45 eV. Calibration mix A was infused throughout and masses were automatically detected and corrected using a 100 ppm window and minimum height of 1×10^3^ counts to ensure 5 ppm mass accuracy, with *m/z* 121.050873 and 922.009798 used as reference ions. AMP-derivatised fatty acids as [M+H]^+^ were aligned using Mass Profinder v10.0 (Agilent Technologies) with an RT tolerance of 0.5 min and MS1 tolerance of 10 ppm. Lipids were identified using a custom personal compound database and library (PCDL) generated from the RT of external quality control samples (Table S3) and MS/MS spectra were manually inspected to ensure correct MUFA isomer identification. Using this method, FA 18:1 isomers were only partially separated and so these data were not analysed further. Data were exported, blank subtracted and quantified from internal standards before being normalised to total protein.

### Knockdown by small interfering RNA

Knockdown experiments were performed in adherent cells grown in 6 well plates for 72 h. Cells were seeded at 1-3×10^5^ cells/well and allowed to adhere overnight. Cells were transfected with Silencer Select siRNA targeting *ACACA* (s883), *FASN* (s5030), *SCD* (s12505,s12504) *FADS2* (s18025,s18023) or negative control (Silencer Select Negative Control No. 1) using Lipofectamine RNAiMAX transfection reagent as per the manufacturer’s instructions (ThermoFisher Scientific).

### RNAseq dataset analysis

uPDAC tumour RNAseq data previously published by Zhou et al. ^41^ were used in this analysis. Genes of interest (*ACACA*, *FASN*, *SCD*, *FADS2*) were filtered from the main dataset, alongside patients who received either neo-adjuvant (Neoadj, n=9) or adjuvant (chemo-naïve, n=66) gemcitabine/nab-paclitaxel treatment. Normalised counts were compared between the groups by unpaired t-test (p < 0.05).

### Statistical analysis

Unless otherwise stated, all data were analysed by comparing CON with GEMR or CombAT cell lines using an unpaired t-test with statistical significance set at *P* < 0.05. Data are expressed as means ± standard error of the mean (SEM).

## Abbreviations

ACC: acetyl-CoA carboxylase
AMP^+^: *N-*(4-aminomethylphenyl)pyridinium
BSA: bovine serum albumin
CE: cholesteryl ester
Cer: ceramide
CL: cardiolipin
CombAT: combination therapy attenuated
CON: control
DG: diacylglycerol
ELOVL: elongation of very long chain fatty acid protein
FADS2: fatty acid desaturase 2
FAS: fatty acid synthase
FOLFIRINOX: folinic acid, 5-fluorouracil, irinotecan and oxaliplatin
GEMC: gemcitabine
GEMR: gemcitabine resistant
IC_50_: half maximal inhibitory concentration
LC-MS: liquid chromatography-mass spectrometry
LPC: lysophosphatidylcholine
LPE: lysophosphatidylethanolamine
MUFA: monounsaturated fatty acid
OzFAD: ozone-induced fatty acid discovery
PBS: phosphate buffered saline
PC: phosphatidylcholine
PDAC: pancreatic ductal adenocarcinoma
PE: phosphatidylethanolamine
PG: phosphatidylglycerol
PS: phosphatidylserine
PTX: paclitaxel
PUFA: polyunsaturated fatty acid
SCD1: stearoyl-CoA desaturase 1
SFA: saturated fatty acid
siRNA: small interfering siRNA
SM: sphingomyelin
TBST: Tris-buffered saline with 0.1% Tween20
TG: triacylglycerol
VEH: vehicle

## Acknowledgements

The authors acknowledge the use of the UNSW Sydney Mark Wainwright Bioanalytical Mass Spectrometry Facility, QUT Central Analytical Research Facility, and Victor Chang Cardiac Research Institute Innovation Centre Metabolomics & Microimaging Facilities (funded by the New South Wales Government Ministry of Health) in this work. The authors also acknowledge and are grateful for input on the project from cancer consumers Louise Bailey & Sundus Javed.

## Funding sources

This project was supported by a Young Investigator Priority-driven Collaborative Cancer Research Grant (APP1160909, funded by Cure Cancer Australia and Can Too!) awarded to SEH, a Miltenyi Research Award (2022) & Victor Chang Cardiac Research Institute Outstanding Early and Mid-career Researcher Grant award to OC, and a PanKind Accelerator grant awarded to NT, SEH, and SJB. BLJP and SJB acknowledge support for enabling technologies from the Australian Research Council Discovery (DP190101486) and Linkage Project (LP180100238).

## Data availability

Analysis and raw files are available for all mass spectrometry data on Metabolomics Workbench, including untargeted lipidomics for Panc1 (ST003757) GEMR, MiaPaCa2 (ST003758) GEMR, & both CombAT cell lines (ST003760); fatty acids analysis for uniformly 13C palmitate-labelled cells (ST003762), CombAT cells (ST003761) & siRNA knockdown (ST003763); and OzFAD data for GEMR cell lines (ST003764).

## Conflicts of Interest

The authors have no conflict of interest to declare.

## Notes

### Competing Interest Statement

The authors have declared no competing interest.

### Summary of Updates

We have added in study IDs for all data files uploaded to Metabolomics Workbench under "Data availability"

