## Supplementary Material for "Targeting *de novo* lipogenesis improves gemcitabine efficacy in pancreatic ductal adenocarcinoma"

### Supplemental methods

#### Measurement of cell proliferation

Cell proliferation was measured by seeding cells at $3\times10^4$ cells per well in 12-well plates. Cells were allowed to adhere overnight, and cell culture media was replaced on days 0 and 3. On indicated days serial plates were stained with crystal violet. Cells were washed once with PBS before being fixed in a solution of 0.5% w/v crystal violet in 50% v/v methanol for a minimum of 10 min. After fixation, cells were washed with water to remove excess crystal violet and then dried overnight. Once dry, wells were solubilised in 1% w/v SDS by vigorous shaking over several hours and the absorbance was read at 570 nm. Relative cell growth was determined by normalising absorbance values to that obtained on day 0. Curves were fitted using an exponential growth equation using least squares regression. Fitted curves were compared using the extra sum-of-squares F test with statistical significance set at $p = 0.05$.

#### Palmitate uptake and oxidation measurement

For palmitate uptake and oxidation measurement cells were grown to near confluency in a 6 well plate. Cells were washed with PBS before the addition of DMEM containing 1 g/L D-glucose and bovine serum albumin (BSA)-conjugated palmitate

For palmitate uptake, 1 mL media containing 200 µM unlabelled palmitate and 2 µCi/mL [1-14C]-palmitate (Perkin Elmer, Sydney, NSW, Australia) conjugated to 2% BSA was added to adherent cells and cells were incubated for 4 h at 37°C. Following this, cells were washed, scraped into ice-cold PBS, and lysed by repeatedly passing through a fine gauge needle. An aliquot was taken for protein determination by BCA assay. Lipids were extracted from the remaining sample and radiolabelled content was counted using a beta counter as described in the methods section of the manuscript.

For palmitate oxidation, cells were incubated in 1 mL of media containing 100 µM palmitate and 0.5 µCi/mL [1-14]-palmitate (conjugated to 2% BSA). Media was removed from samples and added to 1 M perchloric acid to release $CO_2$. The $CO_2$ was captured in 1 M NaOH over 2 h, after which it was counted using a beta counter as described in methods section of the manuscript. The acidified media was collected, centrifuged, and radiolabelled acid soluble metabolite content was measured from the supernatant using a beta counter as described in the methods section of the manuscript. Values measured in both the $CO_2$ and acid soluble metabolite fractions were combined to give total palmitate oxidation. Data were normalised by total protein using a BCA assay quantified from lysed cell pellets as described above.

#### Supplemental data

#### Supplementary Figures

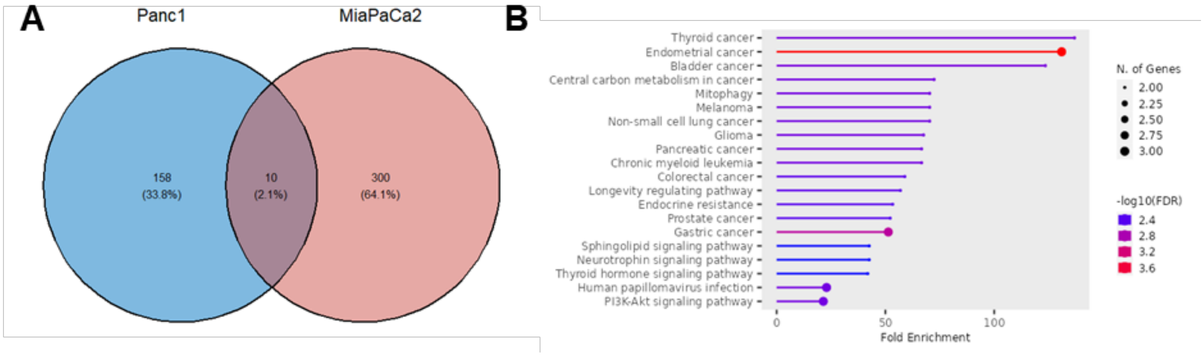


**Figure S1:** Mutational landscape of Panc1 and MiaPaca2 PDAC cells. **A** Venn diagram showing the 10 common genetic mutations between both cell lines (data source: Depmap CCLE). The specific mutations present in these common genes are listed in Table S1. **B** GO enrichment for commonly mutated genes confirms no specific pathway enrichment for lipid metabolism in the top 10 enriched pathways. GO enrichment performed using ShinyGO v0.77 (Ge SX, Jung D & Yao R 2020, *Bioinformatics* 36:2628–2629)

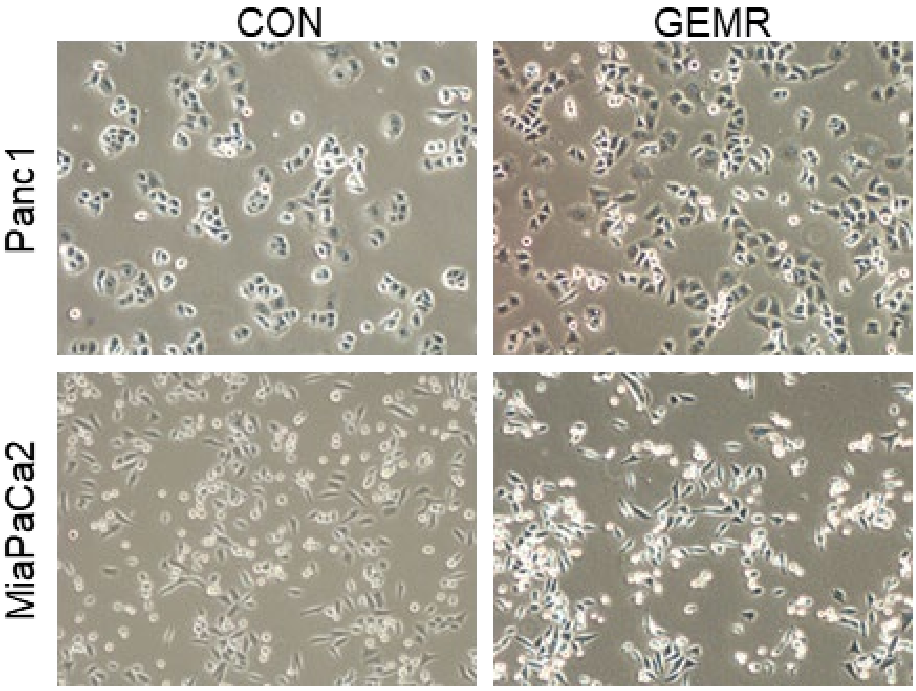


**Figure S2:** light microscopy of both control and GEMR Panc1 & MiaPaCa2 cells at 100x magnification.

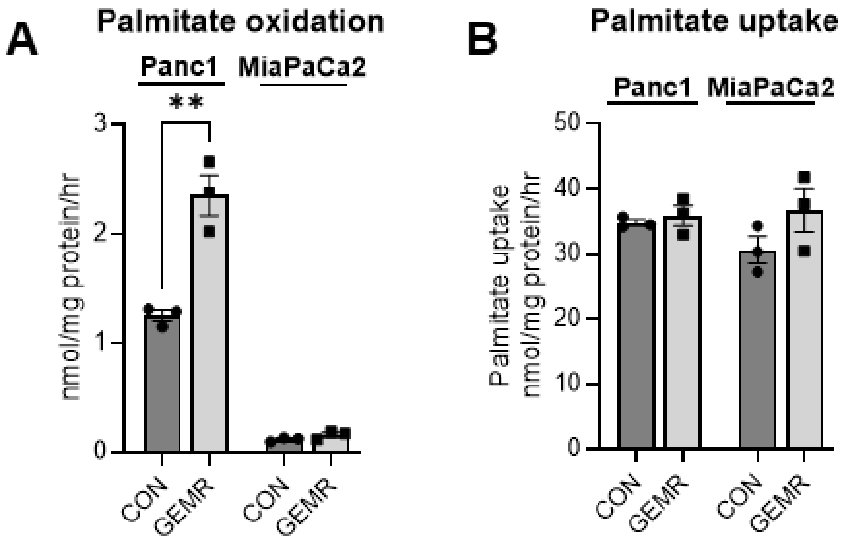


**Figure S3:** measurement of palmitate **A** oxidation and **B** uptake in Panc1 & MiaPaCa2 control (CON) and gemcitabine-resistant (GEMR) cell lines using radiolabelled palmitate. Unpaired t-test * $P<0.05$, ** $P<0.01$, *** $P<0.001$.

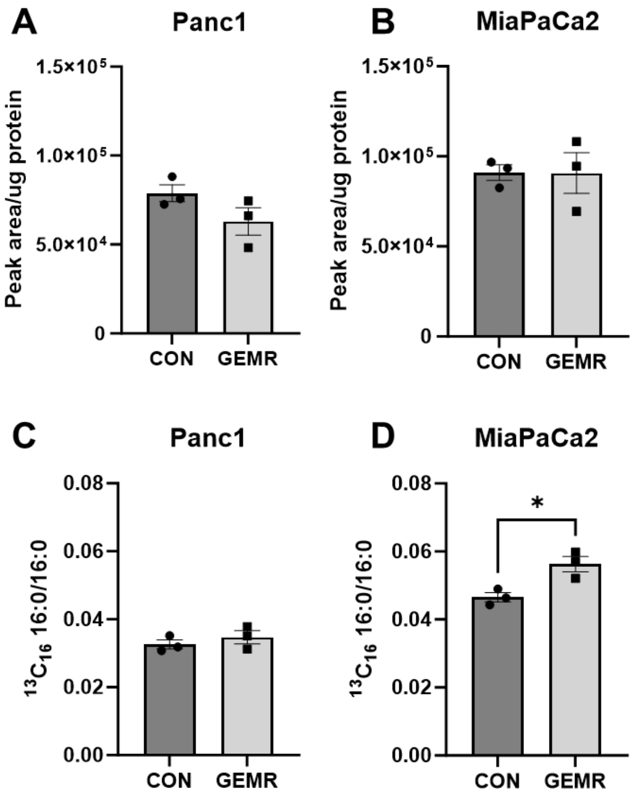


**Figure S4:** Relative amount of palmitate (16:0) in **A** Panc1 and **B** MiaPaCa2 control (CON) and gemcitabine resistance (GEMR) cells following 72 h incubation with $^{13}C_{16}$ palmitate (12.5 μM conjugated to 2% fatty acid free bovine serum albumin). Relative amount of labelled to unlabelled palmitate in the same **C** Panc1 and **D** MiaPaCa2 CON and GEMR cells. Unpaired t test, * $P < 0.05$.

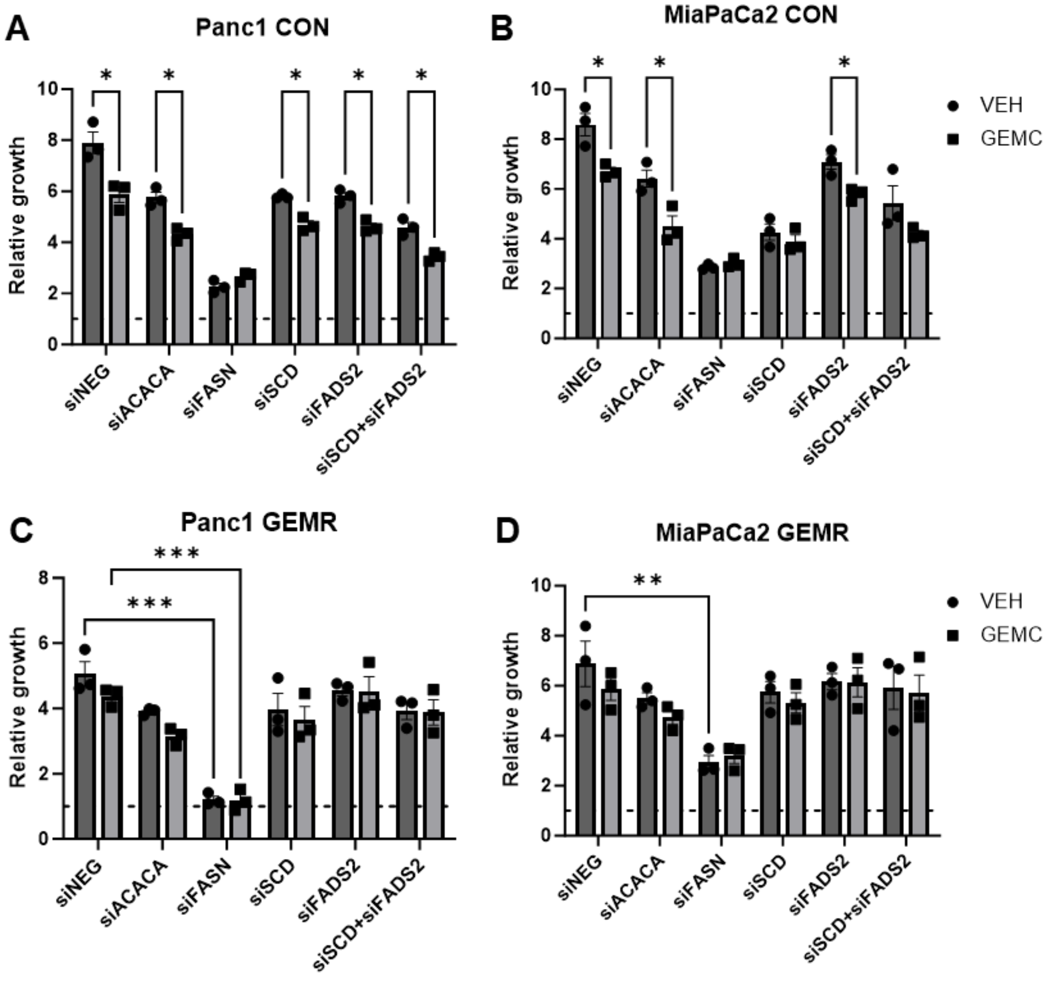


**Figure S5:** effect of 48 h vehicle (VEH) or gemcitabine (GEMC) treatment following 72 h knockdown of lipid synthesis enzymes in **A** Panc1 control (CON) **B** MiaPaCa2 CON **C** Panc1 gemcitabine-resistant (GEMR) or **D** MiaPaCa2 GEMR cell lines. Panc1 CON and GEMR cell lines were treated with 1 μM GEMC while MiaPaCa2 were treated with 100 nM GEMC. Statistical significance was determined by either **A&B** unpaired t-test or **C&D** two-way ANOVA with selected Tukey posthoc comparisons. * $P < 0.05$ ** $P < 0.01$ *** $P < 0.001$. *NEG* negative control *ACACA* acetyl-CoA carboxylase 1 *FADS2* fatty acid desaturase 2 *FASN* fatty acid synthase *SCD* stearoyl-CoA desaturase 1.

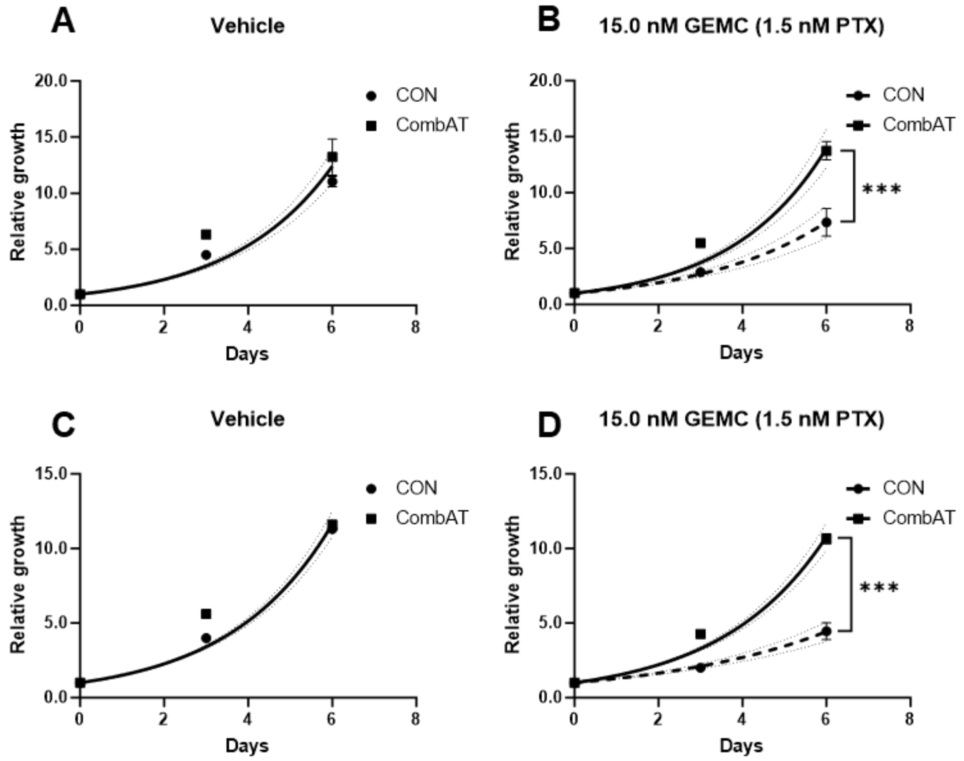


Figure S6: the proliferation of A-B Panc1 and C-D MiaPaCa2 cells in media containing vehicle (VEH) or 15 nM gemcitabine (GEMC)/1.5 nM paclitaxel (PTX). Values are mean±SEM (n=3), *** $P < 0.001$

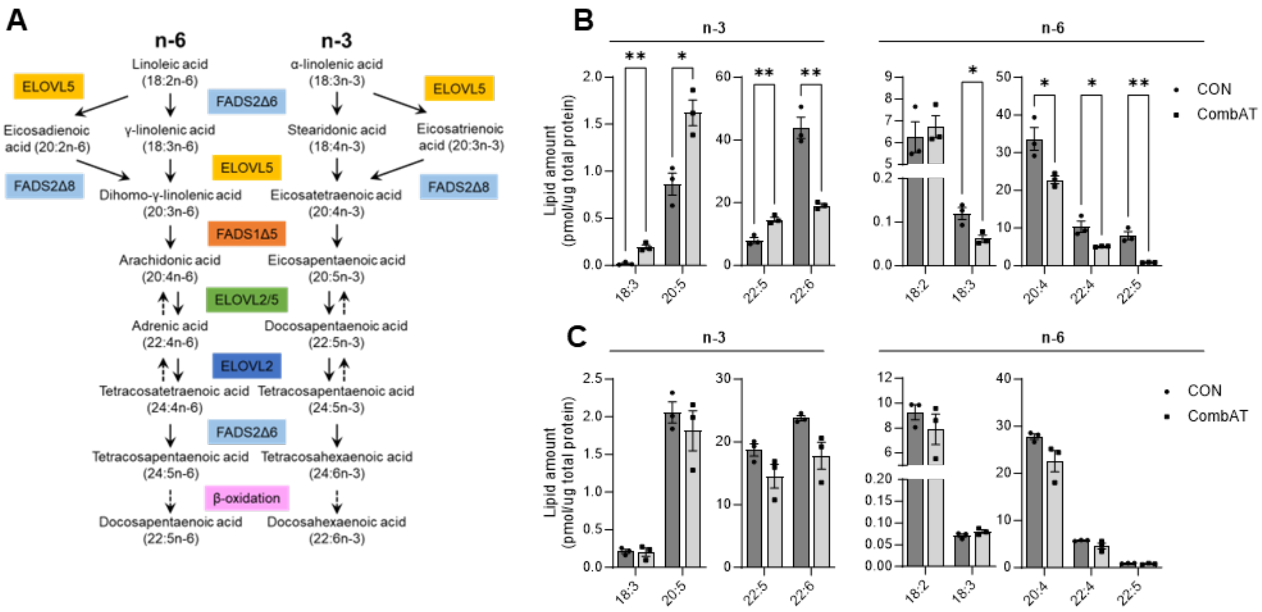


**Figure S7:** **A** pathway of n-3 and n-6 polyunsaturated fatty acids (PUFA) synthesis in mammalian cells. Solid arrows indicated synthesis, dashed indicate β-oxidation. Measurement of $AMP^{+}$-derivatised PUFA in **B** Panc1 and **C** MiaPaCa2 control (CON) and combination therapy attenuated (CombAT) cells. An unpaired t-test determined statistical significance. * $P < 0.05$ ** $P < 0.01$

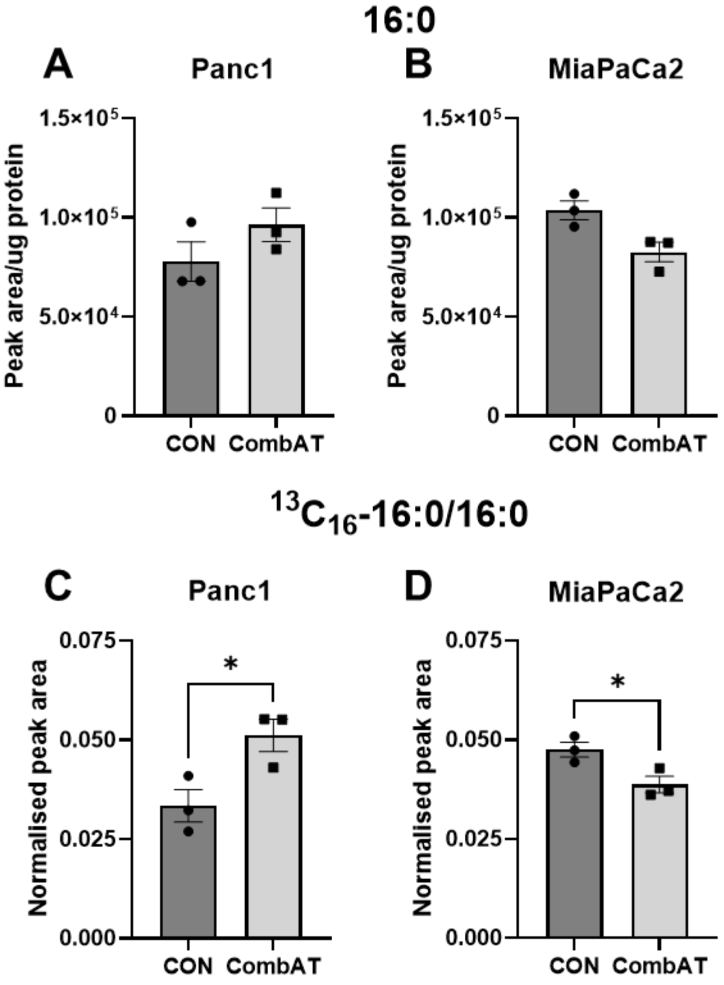


**Figure S8:** Relative amount of palmitate (16:0) in **A** Panc1 and **B** MiaPaCa2 control (CON) and combination therapy attenuated (CombAT) cells following 72 h incubation with $^{13}C_{16}$ palmitate (12.5 µM conjugated to 2% fatty acid free bovine serum albumin). Relative amount of labelled to unlabelled palmitate and labelled 18:0 to labelled 16:0 in the same **C** Panc1 and **D** MiaPaCa2 CON & CombAT cells. Unpaired t test, * $P < 0.05$ ** $P < 0.01$

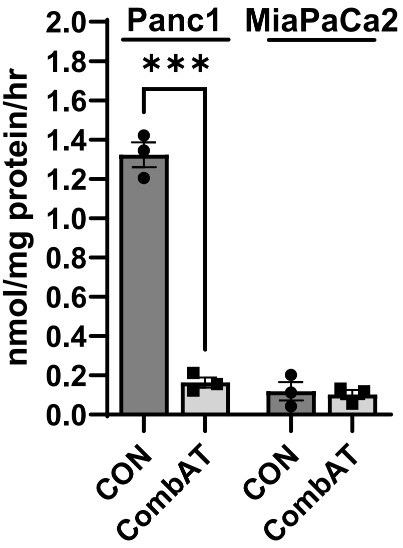


**Figure S9:** Measurement of palmitate oxidation in Panc1 & MiaPaCa2 control (CON) and combination therapy attenuated (CombAT) cell lines. Unpaired t test, *** $P < 0.001$.

#### Supplementary Tables

**Table S1:** Protein mutations in commonly mutated genes of Panc1 and MiaPaCa2 cells

| | Mutation (protein change) | | | |
|---|---|---|---|---|
| Gene | Panc1 | | MiaPaCa2 | |
| | Gene | Type | Gene | Type |
| KRAS | G12D | Missense | G12C | Missense |
| TP53 | R273H | Missense | R248W | Missense |
| BAZ2B | A437T | Missense | G32A | Missense |
| RESF1 | I451M | Missense | S487F | Missense |
| ADGRV1 | W2272R | Missense | R3494S | Missense |
| RELN | C700S | Missense | M709V | Missense |
| CTNNA2 | R538Q | Missense | A207T | Missense |
| SCN9A | T1685I | Missense | E1358K | Missense |
| MGAM | M1553Cfs*36 | FS del | F1771V | Missense |
| NEB | R3616W | Missense | K1882R | Missense |

**Table S2:** Primary antibodies used for immunoblotting experiments.

| **Antibody** | **Catalog #** | **Manufacturer** | **Secondary antibody*** |
|---|---|---|---|
| Phospho-ACC | 3661 | CST | Rabbit |
| Total ACC | 3662 | CST | Rabbit |
| β-actin | sc47778 | SC | Mouse |
| ELOVL2 | A17712 | Abclonal | Rabbit |
| ELOVL5 | A16597 | Abclonal | Rabbit |
| ELOVL6 | A21094 | Abclonal | Rabbit |
| FADS2 | ab72189 | Abcam | Rabbit |
| FAS | 3180 | CST | Rabbit |
| SCD1 | 2438 | CST | Rabbit |

* Secondary horse radish peroxidase-conjugated antibodies were purchased from CST (Mouse #7076, rabbit #7074). *ACC* acetyl-CoA carboxylase, *CST* Cell Signaling Technology (Danvers, MA, USA), *ELOVL* elongation of very long chain fatty acids protein, *FADS2* fatty acid desaturase 2, *FAS* fatty acid synthase, *SC* Santa Cruz Biotechnology (Dallas, TX, USA) *SCD1* stearoyl-CoA desaturase 1.

**Table S3:** AMP-fatty acid retention times for isomer identification by LC-MS/MS.

| Precursor ion mass (m/z) | Fatty acid | RT for identification (min) |
|---|---|---|
| 395.3057 | 14:0 | 2.4 |
| 407.3057 | 15:1 | 2.1 |
| 409.3213 | 15:0 | 3.2 |
| 421.3213 | 16:1n-7,cis | 2.8 |
| 421.3213 | 16:1n-9,cis | 2.9 |
| 421.3213 | 16:1n-10,cis | 3.1 |
| 423.337 | 16:0 | 4.3 |
| 435.337 | 17:1 | 3.6 |
| 437.3526 | 17:0 | 5.5 |
| 445.3215 | 18:3n-3 | 2.45 |
| 445.3215 | 18:3n-6 | 2.6 |
| 447.337 | 18:2n-6,cis | 3.4 |
| 447.337 | 18:2n-6,trans | 3.9 |
| 449.3526* | 18:1n-7,cis | 4.7 |
| 449.3526* | 18:1n-9,cis | 4.8 |
| 449.3526* | 18:1n-10,cis | 4.9 |
| 449.3526* | 18:1n-9,trans | 5.2 |
| 451.3683 | 18:0 | 7.4 |
| 463.3683 | 19:1 | 6.1 |
| 465.3839 | 19:0 | 8.4 |
| 469.3213 | 20:5 | 2.6 |
| 471.337 | 20:4 | 3.4 |
| 473.3526 | 20:3 | 4.5 |
| 475.3683 | 20:2 | 5.5 |
| 477.3839 | 20:1 | 7.9 |
| 479.3996 | 20:0 | 8.7 |
| 493.4152 | 21:0 | 8.9 |
| 495.337 | 22:6 | 3.4 |
| 497.3526 | 22:5n-3,cis | 3.9 |
| 497.3526 | 22:5n-6,cis | 4.5 |
| 499.3683 | 22:4 | 5.1 |
| 503.3996 | 22:2 | 8.2 |
| 505.4152 | 22:1 | 8.7 |
| 507.4309 | 22:0 | 9.1 |
| 521.4465 | 23:0 | 9.4 |
| 533.4465 | 24:1 | 9.1 |
| 535.4622 | 24:0 | 9.5 |

Precursor ions were isolated at a width of 1.3 Th. *RT* retention time
* indicates unresolved isomers with substantial overlaps in retention time, precluding the identification of 18:1 isomers.
